# Phytohormones in *Kappaphycus alvarezii:* Genomic Insights, Activation Thresholds, and Implications for Seaweed-derived Biostimulants

**DOI:** 10.1101/2025.09.19.677482

**Authors:** Bea A. Crisostomo, Leah Ferger, April Mae Tabonda-Nabor, Alexandros Armaos, Gabriel Schweizer, Sergey Nuzhdin, Arturo O. Lluisma, Michael Y. Roleda, Scott Fahrenkrug

## Abstract

The use of seaweed extracts in agriculture has gained significant attention in recent years, yet the genomic mechanisms underlying their activity remain poorly understood. Seaweeds such as *Kappaphycus alvarezii* synthesize diverse bioactive compounds, but their endogenous hormone pathways have been largely uncharacterized. Phytohormones, essential regulators of plant growth, reproduction, and stress responses, are central to the beneficial effects of these biostimulants. To uncover these mechanisms, we performed a genome-wide analysis of hormone biosynthesis and recognition pathways across thirteen algal species, focusing on nine classes of phytohormones. By integrating KEGG annotation, structural modeling, and eggNOG-based orthology, we assessed the completeness and conservation of hormone-associated pathways. Our results revealed consistent retention of key biosynthetic entry points, including IAM-related enzymes for auxin, TRIT1/CYP735A for cytokinins, GA oxidases for gibberellins, and pchA for salicylic acid, alongside divergent or absent canonical upstream enzymes. In parallel, partial but conserved signaling modules were identified for abscisic acid, ethylene, and other classes. This mosaic pattern, fragmented biosynthesis coupled with selectively retained signaling, suggests that algae employ noncanonical enzymatic routes or microbial complementation to sustain hormone activity. Expression profiling in *K. alvarezii* further revealed light-responsive regulation of auxin, cytokinin, GA, and ABA genes, highlighting their role in environmental adaptation. Together, these findings provide new genomic insights into algal hormone biology, establish the first structural and functional evidence for GA metabolism in red algae, and identify candidate phytohormones likely to contribute to the biostimulant activity of *K. alvarezii* extracts.

## Introduction

Seaweed-based biostimulants have emerged as a promising tool in modern agriculture, offering a range of benefits from increased crop growth to stress resilience (Ashour et al, 2023; Frioni et al, 2018; Stasio et al. 2018; Trejo Valencia et al. 2018). The effectiveness of these biostimulants is derived from their ability to enhance nutrient uptake efficiency in various crops (El Boukhari et al, 2020). Plants treated with seaweed extracts have also been shown to have increased stress tolerance to factors such as drought, salinity, and temperature (H. Liu et al, 2019; Masondo et al, 2018; Santaniello et al, 2017; Sharma et al, 2019; Zou et al, 2019). An additional benefit observed for land-based crops is an effectiveness against common pests and diseases (El-Ansary and Hamouda 2014; Machado et al. 2014; Sultana et al, 2012).

The rich biochemical composition of seaweed biostimulants drives their efficacy. Key active components like polysaccharides such as carrageenan, alginates, and laminarins play crucial roles in stress tolerance and plant growth (El Boukhari et al. 2020; Shukla et al. 2019, 2023; Spinelli et al. 2010). Concentrated components of seaweed including vitamins and minerals such as potassium and magnesium, along with phytohormones, like auxins and gibberellins are present in extracts as well, contributing to the overall health and regulation of growth processes in plants (Crouch et al, 1992; De Saeger et al, 2020; Prasad et al, 2010; Stirk et al, 2009).

Phytohormones specifically play an integral role in plant growth, reproduction and environmental response. Auxins, for example, govern directional growth and patterning by mediating cell elongation and distribution of growth regions (Q. Zhang et al, 2022). Gibberellins and cytokinins promote cell division, elongation, seed germination, and delay senescence (Greenboim-Wainberg et al, 2005). In contrast, abscisic acid (ABA) and ethylene often act as growth inhibitors, regulating seed dormancy, fruit ripening, and tolerance to drought and other stressors (Corbineau et al, 2014; Verelst et al, 2010). Other hormones, such as salicylic acid, brassinosteroids, and jasmonates, modulate defense and environmental adaptation (Alazem & Lin, 2015; Shalit-Kaneh et al, 2019). While these signaling molecules have been extensively studied in land plants, their presence and function in algae remain less understood.

Despite the central role of phytohormones in plant biology, their biosynthesis and signaling pathways remain poorly characterized in macroalgae species. This knowledge gap limits the optimization of seaweed cultivation and breeding practices– especially in the face of climate change, disease outbreaks, and increasing demand for sustainable bioactive products (Mo et al, 2020). This challenge is particularly relevant for *Kappaphycus alvarezii (K. alvarezii)*, a genus of red algae cultivated extensively in tropical and subtropical regions. Best known as a primary source of carrageenan, *K. alvarezii* also shows strong potential as a natural biostimulant (Shukla et al, 2023). While such biostimulant effects may be partly mediated by phytohormone-like compounds, the extent to which *K. alvarezii* encodes complete and functional phytohormone biosynthesis and signaling pathways remains unclear. Bridging this knowledge gap is essential for understanding the molecular basis of its bioactivity and for developing targeted applications.

Genomic and transcriptomic resources allow for the investigation of these pathways at the molecular level, offering a powerful strategy to identify the genes and mechanisms involved. Therefore, to address this gap, we investigated the genomic and transcriptomic evidence for phytohormone biosynthesis and signaling pathways in *K. alvarezii* and other algal species. By mapping orthologous genes, modeling key biosynthetic enzymes, and evaluating pathway completeness across species, we aim to clarify the molecular basis of hormone production, regulation, and response in red macroalgae.

Understanding these mechanisms offers insight into the evolution and diversification of hormone signaling outside of land plants, and establishes a molecular foundation for improving seaweed cultivation through hormone-informed breeding strategies. By linking hormone pathway architecture to reproductive and stress-response potential, this work supports the development of higher-yielding, more resilient cultivars and advances the rational design of seaweed-derived biostimulants for agricultural use.

## Methods

### Gene orthology search

Gene orthologs were identified using the *K. alvarezii* Genome explorer (KaGE; https://kage.forjazul.com), web-based platform for exploring the genome and transcriptome of *K. alvarezii*. KaGE includes annotated genomic data from 13 other algal species, these include taxa from multiple, evolutionarily distant lineages: two unicellular red algae (C*yanidioschyzon merolae* and *Galdieria sulphuraria*), seven red macroalgae (*Chondrus crispus, Gracilariopsis chorda, Gracilaria domingensis, Gracilaria parvispora, Pyropia yezoensis, Porphyra linpurpurea, and Porphyra umbilicalis*), the green algae *Chlamydomonas reinhardtii* and *Ulva compressa*, and the brown algae *Ectocarpus siliculosus* and *Saccharina japonica*. While these groups are distant phylogenetically, their inclusion facilitates broad functional comparisons and possible detection of conserved or convergently evolved gene families relevant to phytohormone biosynthesis and signaling.

Orthologous genes involved in phytohormone biosynthesis and signaling were identified using the Kalv-KEGG Explorer (KaGE) tool, which maps annotated proteins to KEGG Orthology (KO) identifiers and highlights their presence within curated pathway maps. KEGG pathways relevant to this study are listed in Table S1, and the resulting gene matches are detailed in Table S2. In addition to KaGE, ortholog identification was cross-validated using KoFamKOALA for KO assignment and eggNOG-mapper (v2) for functional annotation and orthology inference, applying the default score threshold of 60 for eggNOG hits. Using these complementary tools allowed us to capture orthologs supported by different annotation algorithms and reference databases, increasing confidence in pathway reconstruction and minimizing the risk of missing highly divergent candidates.

For *K. alvarezii*, we conducted an additional targeted search using the *K. alvarezii* Annotation Explorer tool. We queried the database with a curated list of phytohormone-related keywords, aliases, KEGG IDs, PFAM domains, and InterPro accessions (Table S3), which yielded 7703 transcripts corresponding to 3029 genes (Tables S4, S5). To prioritize the most likely candidates, we filtered results based on the Sum-of-Categories (SoC) score, a cumulative metric reflecting annotation support across multiple platforms (BLAST2GO, InterProScan, KofamKOALA, HMMR, Wei2GO, eggNOG, orthDB). Higher SoC scores indicate greater cross-platform support and therefore higher confidence in functional relevance. Because SoC scores are influenced by the specificity of the query terms, careful curation was necessary to minimize false positives. The SoC scores ranged from 0.0128 to 12.0 with a mean of 1.21, and a third quartile (Q3) of 1.81 (Table S5; Fig. S1). Among the individual query classes, KEGG ID had the highest mean SoC at 2.21, but also had the highest spread with an interquartile range (IQR) of 2.21. Due to their high specificity, KEGG IDs served as the reference set for establishing an appropriate SoC score threshold. Although the lowest SoC score observed for the KEGG query class was 0.362, the majority had a score of at least 1 (Fig. S1). To improve reliability, we selected 1.0 as the final SoC threshold. The resulting set of candidate genes is listed in Table S6.

### Gene expression analysis in *K. alvarezii*

To assess the transcriptional activity and responsiveness of phytohormone-related genes to stressful light conditions, we analyzed gene expression under varying light conditions. Publicly available transcriptome data from the Marine Science Institute, University of the Philippines Dillman (denoted as “PubMed:MSI” in KaGE) were used via the ShinyKaGE v2.3 interface of the KaGE suite. This dataset consists of RNA-seq profiles from eight (four green- and four brown-color phenotypes) recently domesticated *K. alvarezii* individuals. The algae were maintained for more than a year in an outdoor land-based nursery under medium light intensity (627 ± 64 µmol photons m⁻² s⁻¹ at peak midday irradiance). Propagules were then subjected to two experimental conditions for 28 days: (1) low light (203 ± 6 µmol photons m⁻² s⁻¹) and (2) high light (910 ± 23 µmol photons m⁻² s⁻¹; Tabonda-Nabor et al, 2025).

For KEGG-pathway based expression analysis, the genes with an absolute log2 fold change (logFC) ≥ 1 and a false discovery rate (FDR) ≤ 0.05 were considered to be differentially expressed. To further explore the patterns of gene regulation, we analyzed the expression of the candidate genes (Table S6) identified using *K. alvarezii* Annotation Explorer tool, clustering them based on shared expression profiles (Table S7; Fig. S10).

By integrating expression profiles and ortholog presence across algal lineages, this analysis offers insight into the conservation and divergence of hormone-related gene regulation. Patterns of light-responsive expression in *K. alvarezii* can be compared to orthologous genes in other red algae, and across the phyla, distinguishing conserved stress-response pathways from those that may be lineage-specific or environmentally induced.

### Structural modeling and docking

To identify *K. alvarezii* proteins potentially involved in gibberellin (GA) biosynthesis, we selected 12 candidates spanning key enzymatic steps in the pathway. This targeted analysis was motivated by the failure to detect high-confidence orthologs for canonical GA biosynthetic genes across most species examined under standard stringency thresholds for KofamKOALA and eggNOG (default score cutoff: 60). Despite this apparent absence, previous studies have reported the presence of gibberellins in various algae, including *K. alvarezii,* suggesting that GA biosynthesis may proceed via divergent or noncanonical enzymes (Prasad et al, 2010; Yalçın et al, 2019).

The five GGPPS-like candidates (KAG.1_29607, KAG.1_7012, KAG.1_15296, KAG.1_7476, KAG.1_1991) were selected based on conserved IspA domain architecture and consistent annotation across multiple functional databases. KAG.1_29607, KAG.1_7012, and KAG.1_15296 are fully annotated members of the IspA superfamily, a hallmark of canonical GGPPS proteins, with domain coverage spanning their full sequence lengths. These candidates were also consistently identified by KEGG ortholog mapping and PFAM domain assignment. C1945_pgp086, a chloroplast-localized GGPPS-like protein with strong sequence matches, was also included in GGPPS alignment and docking analyses. Two additional GGPPS-like candidates (KAG.1_7476 and KAG.1_1991) were included due to their possession of IspA domains with N- or C-terminal extensions, suggesting possible alternative isoforms or functional diversification.

The CPS-like candidates (KAG.1_22236 and KAG.1_23590) were selected based on terpene cyclase domain content, KEGG enzyme identifiers, and homology to plant copalyl diphosphate synthases. Additional prioritization considered similarity to enzymes involved in sterol biosynthesis, a known feature of plant diterpene metabolism. To further explore potential GA inactivation mechanisms, two additional candidates (KAG.1_9057 and KAG.1_21559) were evaluated as putative GA2ox-related enzymes. These proteins were aligned to AlphaFold-generated Arabidopsis thaliana and Streptomyces CPS structures to assess structural similarity, and were subsequently docked with ent-copalyl diphosphate (ent-CDP), a known substrate in upstream GA biosynthesis. These analyses aimed to determine whether these proteins could plausibly bind upstream GA intermediates or reflect structural features relevant to inactive GA biosynthesis steps, including conversion to GA8 and GA34.

To identify cytochrome P450 enzymes involved in mid-pathway oxidation steps, we selected two proteins (KAG.1_30825 and KAG.1_11044) as likely ent-kaurenoic acid oxidases (KAO) based on sequence similarity to CYP88A family enzymes and phylogenetic clustering with KAO references. Two additional proteins (KAG.1_8761 and KAG.1_27638) were identified as likely ent-kaurene oxidases (KO) based on homology to CYP701A and CYP714 subfamilies.

To include late-stage GA biosynthesis steps, we selected two proteins (KAG.1_13545 and KAG.1_14123) containing isoprenoid synthase signatures as well as conserved gibberellin oxidase motifs (G2, G3, and G20), consistent with functional roles analogous to GA20ox, GA3ox, and GA2ox.

Three-dimensional structures of all 12 proteins were predicted using ColabFold (AlphaFold2_mmseqs2 notebook v1.5.5) with default parameters. The top-ranked structural model was selected for downstream analysis. Reference structures for KAO, GGPP synthase, and CPS were obtained from the AlphaFold Protein Structure Database. For KO, a structure was generated from the A. thaliana chloroplast ent-kaurene oxidase FASTA sequence using ColabFold. The GAox reference was retrieved from the RCSB Protein Data Bank (PDB ID: 3EBL).

Pairwise structural alignments were conducted using the RCSB PDB Structure Alignment Tool (Bittrich et al, 2024) with TM-align. Root-mean-square deviation (RMSD), structural overlap, and conservation of catalytic domains were evaluated to assess similarity and functional relevance.

Ligand docking was performed using SwissDock “Attracting Cavities” mode to focus the search on predicted surface pockets (Grosdidier et al, 2011; Zoete et al, 2016; Röhrig et al, 2023; Bugnon et al, 2024). Ligands included key pathway intermediates and products: GGPP, ent-copalyl diphosphate (ent-CDP), ent-kaurene, ent-kaurenoic acid, GA12, GA3, and GA4. Ligands were sourced from PubChem and entered into SwissDock using SMILES sequences. For each protein–ligand pair, binding free energy (ΔG), docking cluster count, and spatial positioning were assessed to evaluate predicted substrate affinity and enzymatic plausibility.

## Results

To explore phytohormone biosynthesis and signaling in *K. alvarezii,* we analyzed candidate genes across nine major hormone classes: auxin, cytokinin, gibberellin, abscisic acid, ethylene, brassinosteroids, jasmonates, salicylic acid, and strigolactones. For each hormone, we examined the orthologous gene presence, pathway completeness, and domain conservation using genomic and protein-level annotation, supplemented by structural modeling and gene expression data. The following subsections present results grouped by hormone class.

### Auxin

Auxins, primarily represented by indole-3-acid (IAA), are synthesized via both tryptophan (Trp)-dependent and Trp-independent pathways. Four Trp-dependent routes have been described in plants: IPyA, IAOx, TAM, and IAM, with the IPyA pathway considered the major route in embryophytes (Casanova-Sáez et al, 2021). Although a Trp-independent pathway has been proposed (Normanly et al, 1993), its mechanism remains unresolved (Nonhebel, 2015).

Auxin biosynthesis in algae appears to diverge from classic plant pathways. Of the species analyzed, the green alga, Ulva compressa encoded an ortholog of tryptophan aminotransferase (TAM1), which catalyzes the conversion of tryptophan (Trp) to indole-3-pyruvic acid (IPyA). In contrast, the downstream enzyme, YUCCA, which converts IPyA to IAA, was detected only in the brown algae, *Saccharina japonica* and *Ectocarpus siliculosus* (Fig. 1). The absence of a complete IPyA pathway in most algal species suggests that, unlike embryophytes, they do not rely on this route for auxin biosynthesis.

**Fig. 1.**
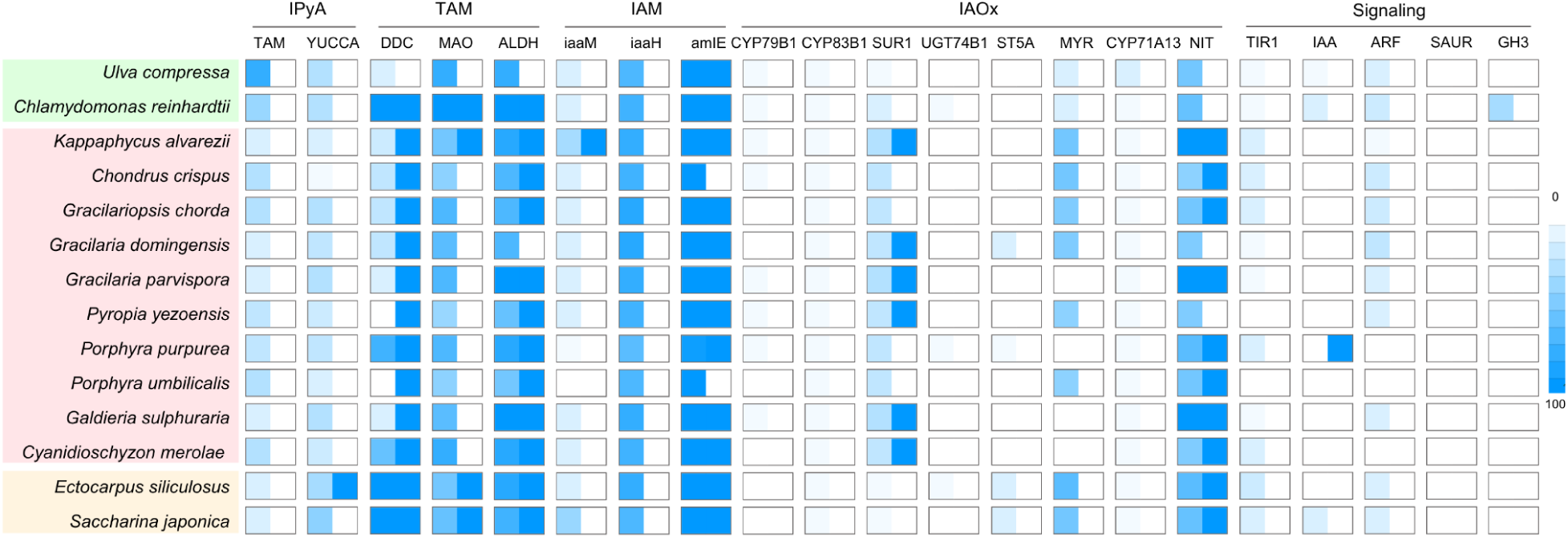
Heatmap comparison of algal orthologs for genes involved in auxin biosynthesis (IPyA, TAM, IAM, IAOx pathways) and signaling (TIR1, ARF, SAUR, GH3). Species are grouped by major algal lineages (chlorophytes, rhodophytes, glaucophytes, brown algae). Color intensity indicates KofamKOALA (left) and EggNOG (right) orthology scores (0–100).

KofamKOALA and EggNOG scores for IPyA pathway genes were generally low or absent across all algal species, further supporting limited conservation of this route. However, the possibility of functionally analogous, distantly related enzymes cannot be excluded.

Partial orthologs of genes in the IAOx pathway were found in several rhodophytes, though CYP79B2, which initiates this pathway in Arabidopsis, was absent from all algal species examined. Consistent with prior reports, this pathway appears specific to Brassicaceae (Casanova-Sáez et al, 2021; Mikkelsen et al, 2000; Sugawara et al, 2009). Annotation scores for detected IAOx-pathway genes were low, reflecting uncertain orthology and possible alternative metabolic functions (Fig. 1).

All species investigated contained orthologs of TAM pathway enzymes, which suggests the possible usage of this route. However, these genes are known to function in multiple metabolic contexts (Parthasarathy et al, 2018), and overexpression of tryptophan decarboxylase (TDC) in Nicotiana tabacum failed to increase IAA levels (Songstad et al, 1990), indicating that this pathway may play a minor role in auxin biosynthesis in land plants. In *K. alvarezii*, TAM pathway genes showed moderate KofamKOALA and EggNOG scores, consistent with their conserved but multifunctional nature.

In contrast, the IAM pathway appears to be the most conserved in algae. Orthologs of indoleacetamide hydrolase (iaaH) and amidase (amiE) were broadly present, and *K. alvarezi*i uniquely encoded iaaM, the microbial enzyme catalyzing Trp to IAM conversion (Fig. 1). These three IAM-pathway genes consistently scored highest in both KofamKOALA and EggNOG annotations among all auxin-related genes, providing strong homology support for this pathway in *K. alvarezii* (Fig. 1; Table S2). This suggests either gene acquisition from microbes or the presence of an analogous endogenous function. Biochemical studies in Ulva fasciata and Dictyota humifusa further support this pathway as the dominant route for auxin production in algae (Stirk et al, 2009).

For transport, only PILS (PIN-LIKES) family orthologs were detected in *K. alvarezii* (Table S6), while AUX1 and canonical PIN proteins were absent. The PILS ortholog in *K. alvarezii* has moderate annotation scores, suggesting a conserved but possibly divergent transport role, while AUX1/PIN orthologs were undetected. Because PILS proteins and ABCB transporters contribute to auxin movement in plants (Geisler et al, 2017), these may mediate auxin transport in algae as well.

Auxin signaling in plants depends on the SCF E3 ligase complex and the TIR1/AFB-Aux/IAA-ARF axis. All algal species encoded core SCF complex components CUL1, SKP1, and E3 ligase), but canonical auxin signaling proteins were absent. KofamKOALA and EggNOG scores were high for CUL1, SKP1, and E3 ligase, but negligible for TIR1, ARF, and Aux/IAA, with only one low-scoring putative TIR detected in *K. alvarezii* (Fig. 1, S2; Table S6).

Despite the genomic gaps, physiological studies have shown that algae respond to exogenous auxin, influencing growth, morphology, and developmental polarity (Garcia-Jimenez et al, 1998; Fries and Āberg, 1978; Basu et al, 2002). Auxin levels are also modulated by environmental cues: in *Klebsormidium nitens*, high light increases IAA production (Serrano-Pérez et al, 2022). Similarly, in *K. alvarezii*, we observed upregulation of auxin biosynthesis and transport genes under high light (Fig. S3; Table S7).

Together, these findings suggest that while canonical auxin biosynthetic and signaling pathways are incomplete in algae, alternative or modified routes may support functional auxin production and perception. The integration of high-confidence annotation scores with light-responsive expression patterns strengthens the case for a functional IAM pathway in *K. alvarezii*. The conservation of key biosynthetic genes, light-responsive expression patterns, and physiological sensitivity to auxin all point to a conserved or co-evolved role for auxin in algal development and environmental response.

### Cytokinin

In plants, cytokinins are adenine-derived hormones that most commonly occur in the isoprenoid forms isopentenyl adenine (iP), trans-zeatin (tZ), and cis-zeatin (cZ) (Li et al, 2021). While tZ is the most bioactive and prevalent in land plants, cZ is more abundant in microalgae (Xingfeng et al, 2018). Their biosynthesis begins with dimethylallyl diphosphate (DMAPP), produced via the methylerythritol phosphate (MEP) pathway in plastids or the mevalonate (MVA) pathway in the cytosol (Kasahara et al, 2004). DMAPP can be converted to iP by adenylate dimethylallyltransferase (IPT), and then to tZ by cytokinin trans-hydroxylase (CYP735A) (Kakimoto, 2001; Takei et al, 2004). Alternatively, MVA-derived DMAPP may be converted to cZ via tRNA dimethylallyltransferase (TRIT1) (Miyawaki et al, 2006).

Orthologs of both CYP735A and TRIT1 were identified across all algal species examined, suggesting the presence of both tZ and cZ biosynthesis. KofamKOALA and EggNOG scores for CYP735A and TRIT1 were consistently high in most species, including *K. alvarezii*. However, IPT orthologs had much lower scores or were absent, this pattern points to a possible divergence or replacement of canonical IPT enzymes, which may alter or constrain IP formation in algae (Fig. 2). Previous work also shows that macroalgae produce a variety of cytokinin derivatives—including aromatic cytokinins and O-glucosides—despite lacking canonical derivatization enzymes found in embryophytes (Stirk et al, 2003; 2009), pointing to divergent enzymatic machinery.

**Fig. 2.**
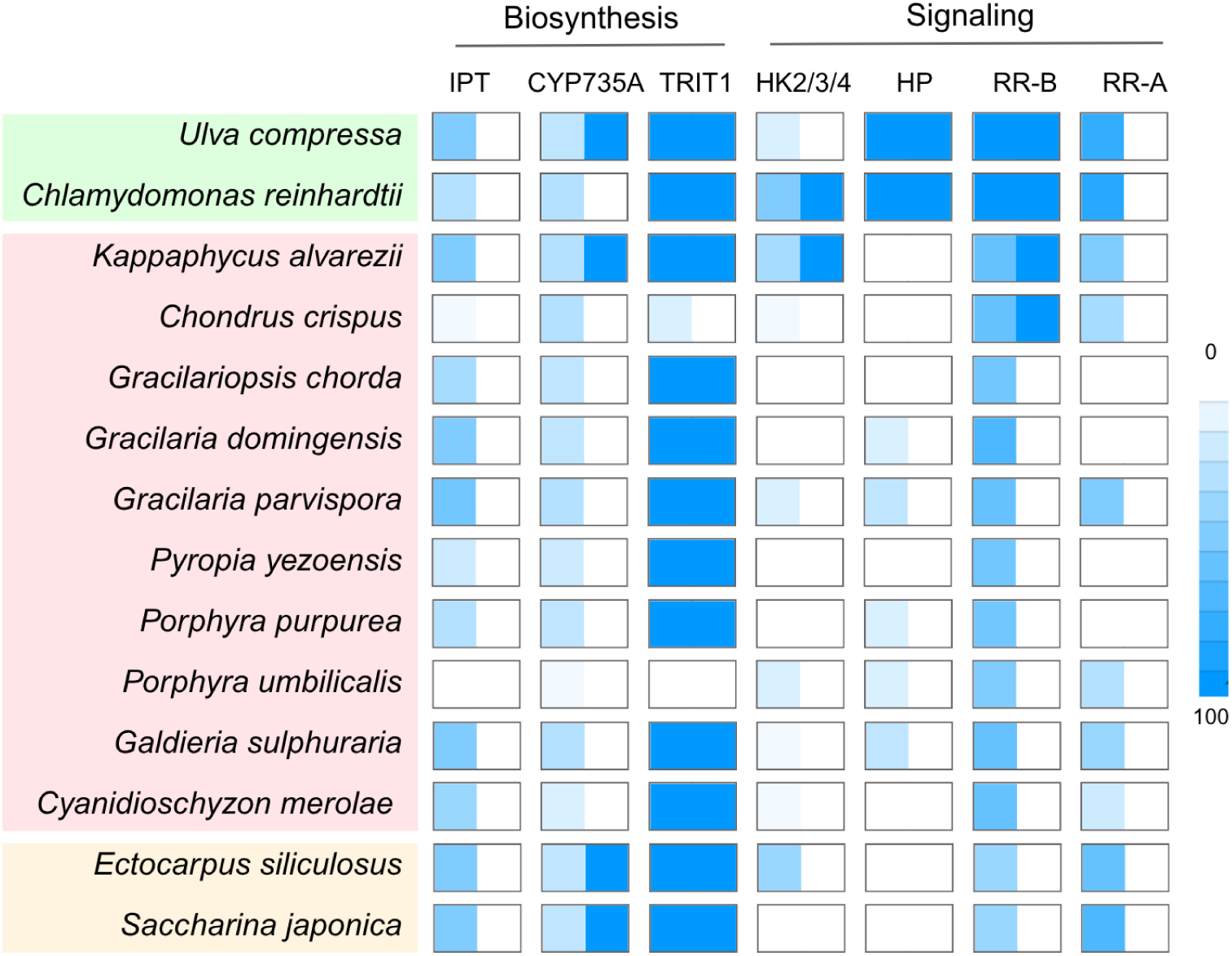
Heatmap comparison of algal orthologs for genes involved in cytokinin biosynthesis (IPT, CYP735A, TRIT1) and signaling (HK2/3/4, HP, RR-B, RR-A). Species are grouped by major algal lineages (chlorophytes, rhodophytes, glaucophytes, brown algae). Color intensity indicates KofamKOALA (left, 0-100%) and EggNOG (right) orthology scores >60).

Cytokinin signaling in plants involves perception by histidine kinase (HK2–4) receptors, which initiate a phosphotransfer cascade through histidine phosphotransfer proteins (HPs) and B-type response regulators (RR-Bs), ultimately activating cytokinin-responsive genes. A-type RRs (RR-As) provide negative feedback regulation. In this study, a complete set of cytokinin signaling genes was found only in green algae. Consistent with the ortholog distribution, KofamKOALA and EggNOG scores for HK2-4 and HPs were zero in phaeophytes, while RR-A and HPs were absent or scored below the threshold in rhodophytes, This reinforces the idea that certain algal groups may have reduced or modified cytokinin signaling capacity, relying on alternative relay systems to mediate cytokinin perception and downstream transcriptional regulation (Fig. 2).

Physiological evidence supports a conserved role for cytokinins in algal growth and reproduction. Like in land plants, cytokinins promote algal growth (Burkiewicz, 1987; Lin & Stekoll, 2007; Araújo et al, 2021) and have been linked to seasonally driven developmental shifts. In Macrocystis pyrifera, cytokinin-like activity peaks in spring and autumn—coinciding with elevated levels of freebase and riboside cytokinins—and declines in summer and winter, when O-glucoside forms dominate (de Nys et al, 1990). Similar seasonal trends are observed in Ecklonia maxima (Featonby-Smith & Van Staden, 1984) and reproductive cycles in Sargassum heterophyllum (Mooney & Van Staden, 1984).

In *K. alvarezii*, the integration of annotation scores and expression analysis provides further detail. High light conditions upregulate genes involved in cZ biosynthesis, while tZ biosynthesis and cytokinin response are enhanced under medium light (Fig. S4; Table S7). High-confidence KofamKOALA and EggNOG annotation scores for cZ-biosynthetic genes align with their strong light-dependent expression, where tZ-pathway genes, despite moderate annotation scores, showed higher expression under medium light. This suggests that regulatory tuning, rather than structural gene loss, may explain the expression differences between the two pathways (Fig. S4). This pattern may reflect light-specific modulation of cytokinin function, as previously observed in Arabidopsis, where cytokinin signaling confers protection against light-induced stress (Cortleven et al, 2014). Additionally, in Zea mays, cZ was found to enhance antioxidant activity (Yousaf et al, 2024), potentially explaining its upregulation under high light in *K. alvarezii*.

Overall, our findings suggest that *K. alvarezii* retains the genetic capacity for cytokinin biosynthesis and displays light-responsive regulation of key cytokinin-related genes. By combining high-confidence annotation scores with expression data, we can infer that these pathways are likely functional despite the absence of some canonical signaling components in red algae. In addition to endogenous biosynthesis, it is also possible that associated microbial communities may contribute to the presence of cytokinin, as many marine bacteria and endophytes are known to produce cytokinins or cytokinin-like compounds (Amin et al, 2015; Singh & Reddy, 2016). Such microbial actions may act to complement or modulate host cytokinin signaling, especially under environmental stress.

### Gibberellin

Gibberellic acid (GA) is synthesized from geranylgeranyl diphosphate (GGPP), a 20-carbon compound formed from isopentyl diphosphate (IPP) and dimethylallyl diphosphate (DMAPP). GGPP undergoes two-step cyclization catalyzed by ent-copalyl diphosphate synthase (CPS) and ent-kaurene synthase (KS), forming ent-kaurene, a tetracyclic diterpene precursor (Sun & Kamiya, 1994). In plants, these reactions are plastid-localized and typically driven by the MEP pathway, with the mevalonic acid (MVA) pathway contributing minor input (Kasahara, 2002). Ent-kaurene is then oxidized by ent-kaurene oxidase (KO) to produce ent-kaurenoic acid, which is further oxidized by ent-kaurenoic acid oxidase (KAO) to form GA12. GA12 serves as a central precursor converted into a variety of bioactive GAs through GA 20-oxidases (GA20ox) and GA 3-oxidases (GA3ox), or inactivated by GA 2-oxidases (GA2ox).

Canonical GA pathway genes were not recovered under standard annotation thresholds. In particular, CPS and KS orthologs were absent or highly divergent in *K. alvarezii.* Domain-level searches, however, identified candidate terpene synthases and cytochrome P450 enzymes spanning both upstream and downstream steps of the pathway (Fig. 3).

**Fig. 3.**
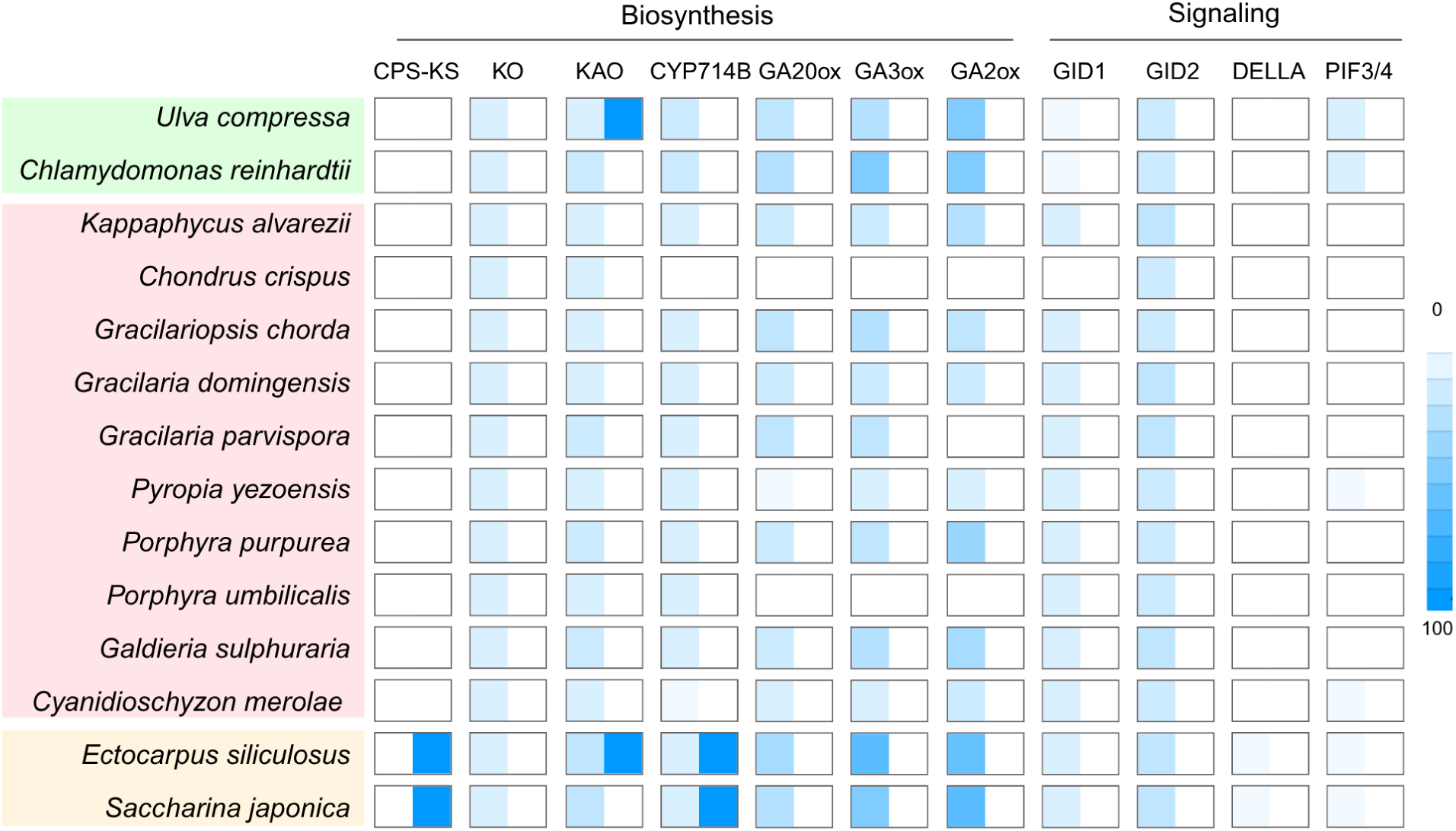
Heatmap comparison of algal orthologs for gibberellin biosynthesis (CPS-KS, KO, KAO, CYP714B, GA20ox, GA3ox, GA2ox) and signaling (GID1, GID2, DELLA, PIF3/4). Species are grouped by major algal lineages (chlorophytes, rhodophytes, glaucophytes, brown algae). Color intensity indicates KofamKOALA (left) and EggNOG (right) orthology scores (0–100) from KofamKOALA and EggNOG.

Structural modeling and alignment supported these identifications. GGPPS-like candidates displayed a range of similarity to *Arabidopsis thaliana* references, with some exhibiting conserved domain features indicative of enzymatic potential (Fig. 4a; Table S8). CPS-like candidates, by contrast, showed consistently poor alignments by both plant and bacterial references, suggesting either functional divergence or absence of canonical CPS activity. Downstream candidates, including putative KAO, KO, and GA20ox proteins retained strong structural homology or conserved catalytic motifs relative to reference enzymes (Fig. 5a; Table S8a).

**Fig. 4.**
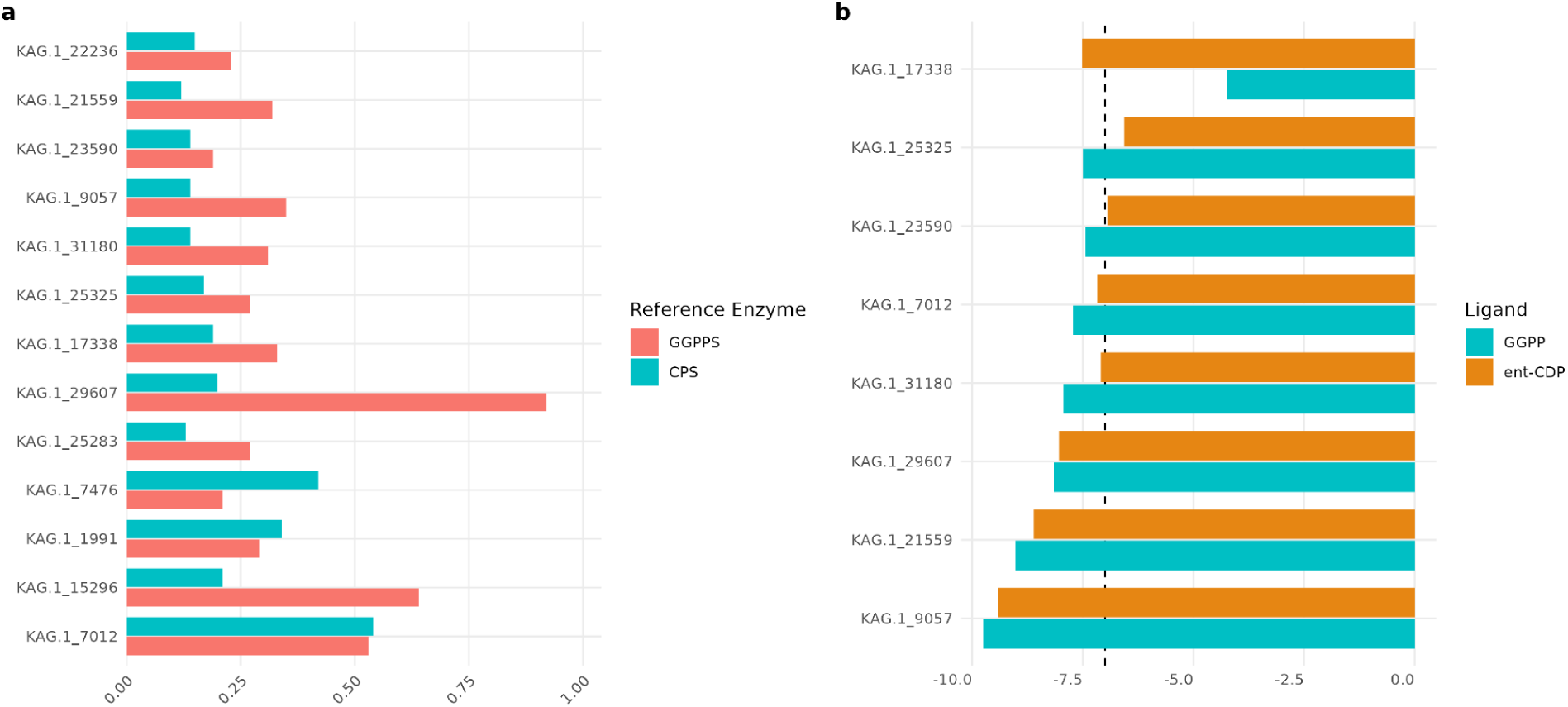
(a) TM-scores from pairwise structural alignments of GGPPS-like (red) and CPS-like (aqua) candidate proteins against *Arabidopsis thaliana* reference enzymes. TM-scores closer to 1 indicate greater structural similarity (b) Predicted binding free energies (ΔG, kcal/mol⁻¹) for GGPP (teal) and ent-CDP (orange) docked to upstream candidates using Swissdock “Attracting Cavities” mode. Lower ΔG values indicate stronger predicted binding affinity.

**Fig. 5.**
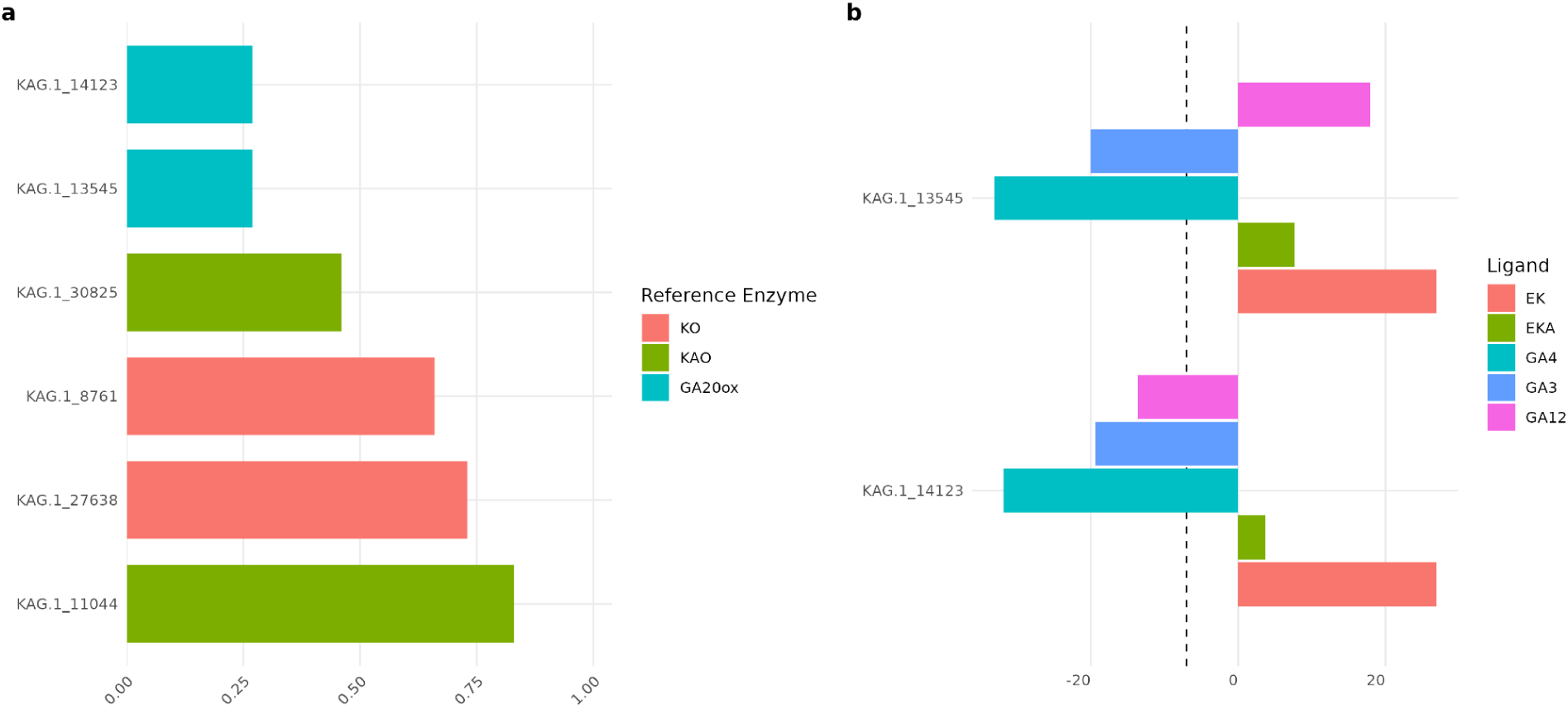
(a) TM-scores from pairwise structural alignment of KO-like, KAO-like, and GA20ox-like candidate proteins against *Arabidopsis thaliana* reference enzymes. TM-scores closer to 1 indicate greater structural similarity (b) Predicted binding free energies (ΔG, kcal/mol⁻¹) for GA intermediates GA3, GA4, GA12, ent-kaurene, and ent-kaurenoic acid docked to top two downstream candidates using Swissdock “Attracting Cavities” mode, lower Δ G values indicate stronger predicted docking affinity

Ligand docking results reinforced these patterns. GGPP docking indicated plausible activity for select GGPPS-like isoforms (Fig. 4b; Table S9). Although CPS-like proteins had low alignment quality, docking revealed potential compatibility with ent-CDP for a subset of candidates. Downstream oxidase candidates demonstrated predicted binding to GA intermediates (GA12, GA3, GA4), with strongest affinity suggested for GA4 (Fig. 5b; Table S9a).

Expression profiling showed that GAox candidates (KAG.1_13545 and KAG.1_14123) were moderately expressed under medium-light conditions, consistent with roles in GA activation. GA2ox-like candidates, including KAG.1_9057 and KAG.1_21559, were strongly upregulated under high-light conditions, aligning with a potential role in GA inactivation. Docking of these GA2ox-like proteins to GA8 and GA34 further supported their structural capacity to catalyse inactivation steps.

Comparative analysis revealed lineage-specific differences: while red algae, including *K. alvarezii* lacked CPS/KS and most KO orthologs, brown algae such as *Ectocarpus* and *Saccharina* retained both upstream and downstream enzymes. In *K. alvarezii*, however, signaling components such as GID2 and PIF3/4 were detected, suggesting that partial GA responsiveness is conserved.

### Abscisic Acid

In plants, abscisic acid (ABA) is synthesized from zeaxanthin via a multi-step carotenoid cleavage pathway. Zeaxanthin is first epoxidized to violaxanthin by zeaxanthin epoxidase (ZEP), then cleaved by 9-cis-epoxycarotenoid dioxygenase (NCED) to produce xanthoxin. Xanthoxin is converted to abscisic aldehyde by ABA2 and finally oxidized to ABA by aldehyde oxidase (AAO3), with ABA3 supplying the molybdenum cofactor (Barrero et al, 2008; Bittner et al, 2001; Seo et al, 2000).

Ortholog searches revealed limited conservation of this pathway in *K. alvarezii*. ZEP and NCED were detected in chlorophyte and phaeophyte transcriptomes, but most downstream enzymes were absent, except ABA3, which was also identified in some red algae (Fig. 6). In *K. alvarezii*, only ABA3 was recovered from the biosynthetic pathway, with no evidence for ZEP, NCED, ABA2, or AAO3.

**Fig. 6.**
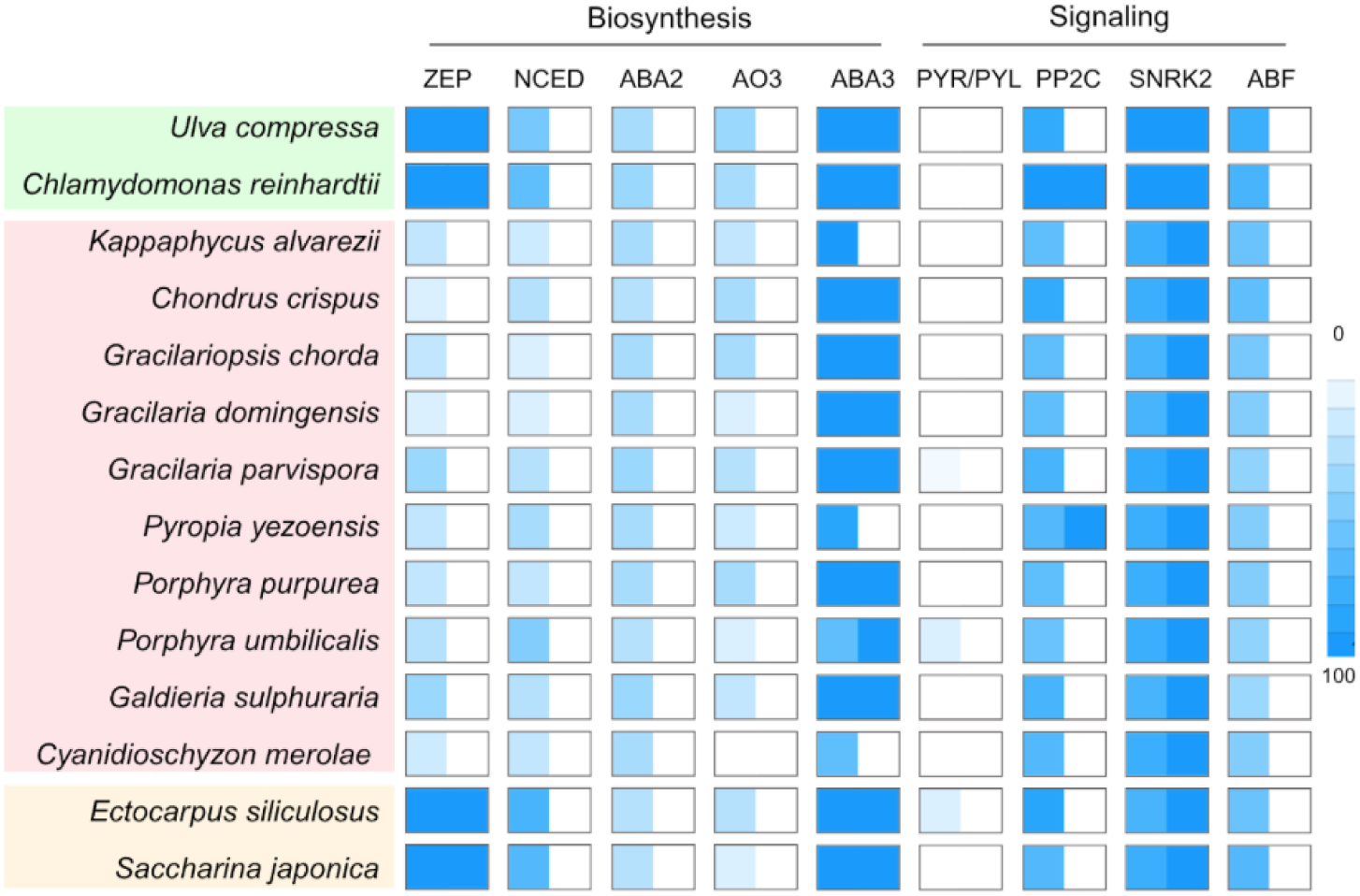
Heatmap comparison of algal orthologs for abscisic acid (ABA) biosynthesis (ZEP, NCED, ABA2, AO3, ABA3) and signaling (PYR/PYL, PP2C, SNRK2, ABF). Species are grouped by major algal lineages (chlorophytes, rhodophytes, glaucophytes, brown algae). Color intensity indicates KofamKOALA (left 0-100%) and EggNOG (right, >60) orthology scores.

In contrast, signaling pathway components were better represented. Orthologs of PP2C, SnRK2, and ABF were detected with strong KofamKOALA and EggNOG scores, while canonical PYR/PYL receptors were absent (Fig. 6).

Expression analysis showed that the synthesis of the ABA precursor, zeaxanthin, was highest in medium light and lowest in low light (Fig. S5). Conversely, the signaling genes, SnRK2 and ABF, peaked under low light, declined under medium light, and displayed mixed responses under high light, suggesting a photoregulatory pattern across biosynthesis and signaling components (Fig. S9).

### Ethylene

Ethylene is synthesized by methothionine via the intermediate S-adenosylmethionine (SAM). SAM is converted to 1-aminocyclopropane-1-carboxylic acid (ACC) by ACC synthase (ACS), and then to ethylene by ACC oxidase (ACO), with ACS acting as the rate-limiting enzyme (Xu et al, 2014). Among all examined algal species, *K. alvarezii* possessed strong ortholog signals for MAT and ACS, but not detectable ACO (Fig. 7). The high KofamKOALA and EggNOG scores for ACS indicate strong annotation support, consistent with a functional capacity for ACC biosynthesis, even in the absence of a canonical ACO ortholog. Prior studies confirm ethylene biosynthesis in seaweeds via the ACC pathway (Broadgate et al, 2004; Plettner et al, 2005; Garcia-Jimenez et al, 2013), suggesting that even with partial pathway components, ethylene production is functional.

**Fig. 7.**
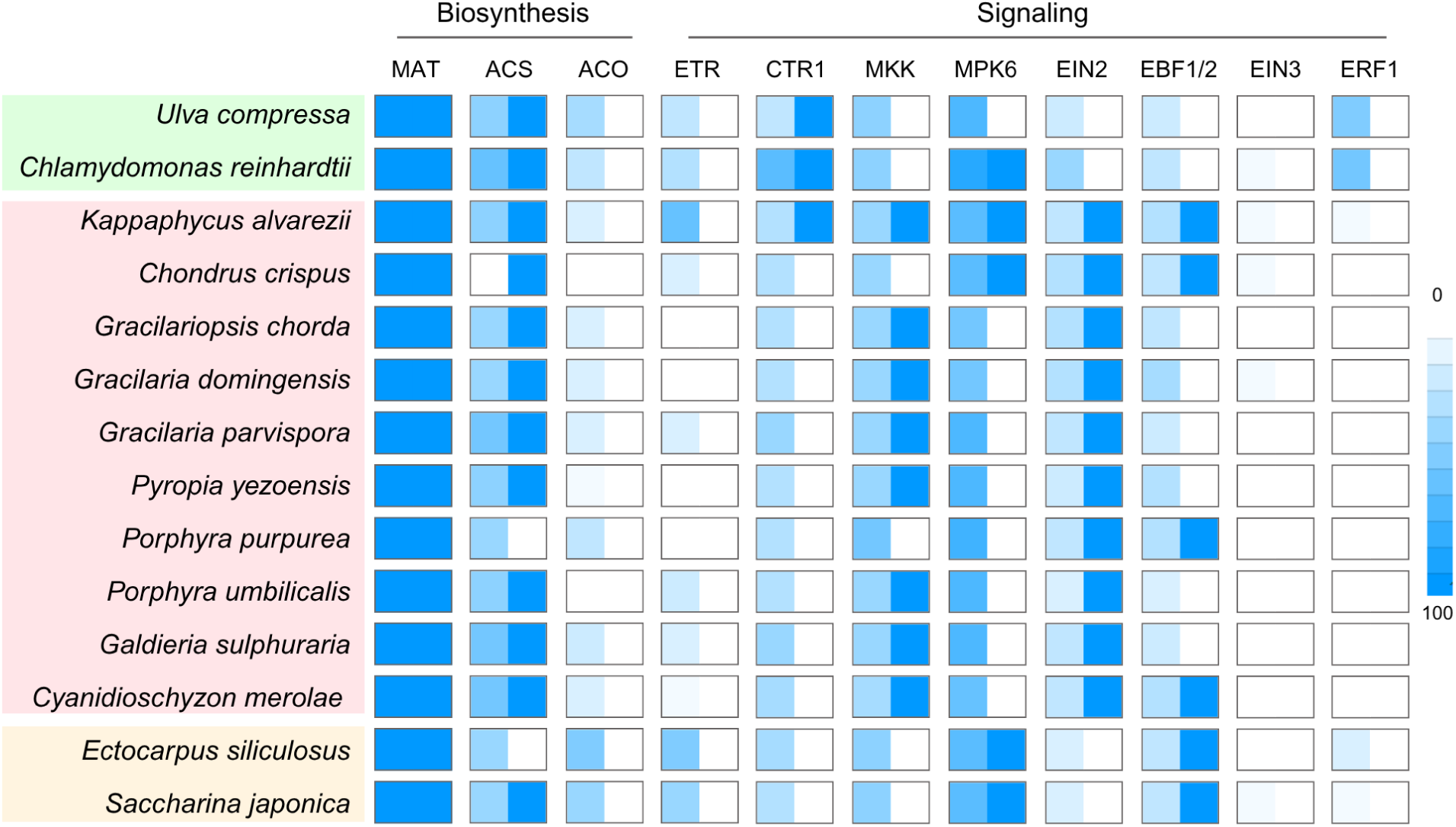
Heatmap comparison of algal orthologs for ethylene biosynthesis (MAT, ACS, ACO, ETR) and signaling (CTR1, MKK, MPK6, EIN2, EBF1/2, EIN3, ERF1). Species are grouped by major algal lineages (chlorophytes, rhodophytes, glaucophytes, brown algae). Color intensity indicates KofamKOALA (left) and EggNOG (right) orthology scores (0–100).

Ethylene perception in land plants involves copper-dependent receptors (ETRs) located in the endoplasmic reticulum. In the absence of ethylene, ETRs activate the kinase CTR1, which phosphorylates and suppresses EIN2, blocking downstream signaling. Ethylene binding inactivates CTR1, enabling EIN2 to translocate to the nucleus, where it stabilizes the transcription factor EIN3 by inhibiting its degradation via EIN3-binding F-box proteins (EBFs).

None of the algae examined possessed a complete ethylene signaling cascade. Chlorophytes lacked the most components, whereas rhodophytes retained all core elements except for EIN3 and ERF1. This suggests that while ethylene signaling may be incomplete, partial pathways may still support perception, and therefore, downstream responses. In *K. alvarezii*, orthologs for CTR1, MKK, EIN2, and EBF1/2 were identified with moderate to strong annotation scores, but EIN3 and ERF1 were absent (Fig. 7). These findings support the possibility of truncated signaling pathways that may still transduce partial ethylene responses.

Expression profiling revealed that ACS was consistently upregulated in high light relative to low light, supporting a light-induced increase in ACC production (Fig. S6). CTR1, the negative regulator of ethylene signaling, also showed elevated expression under high light conditions, suggesting feedback moderation of downstream signaling despite increased precursor synthesis. MPK6, a positive signaling component, was most strongly expressed under medium light conditions, indicating a condition-specific activation of signaling nodes (Fig. S9). In medium light, both positive (MPK6, EIN2) and negative (CTR1) regulators were upregulated, reflecting a balanced modulation of ethylene pathway activity.

### Tryptophan Metabolites

Tryptophan serves as a precursor for several signaling metabolites, including melatonin, serotonin, and kynurenine. Among these, kynurenine and its derivatives appear particularly relevant in *K. alvarezii* due to consistent gene presence and strong expression patterns across light regimes (Fig. 8, Fig. S3).

**Fig. 8.**
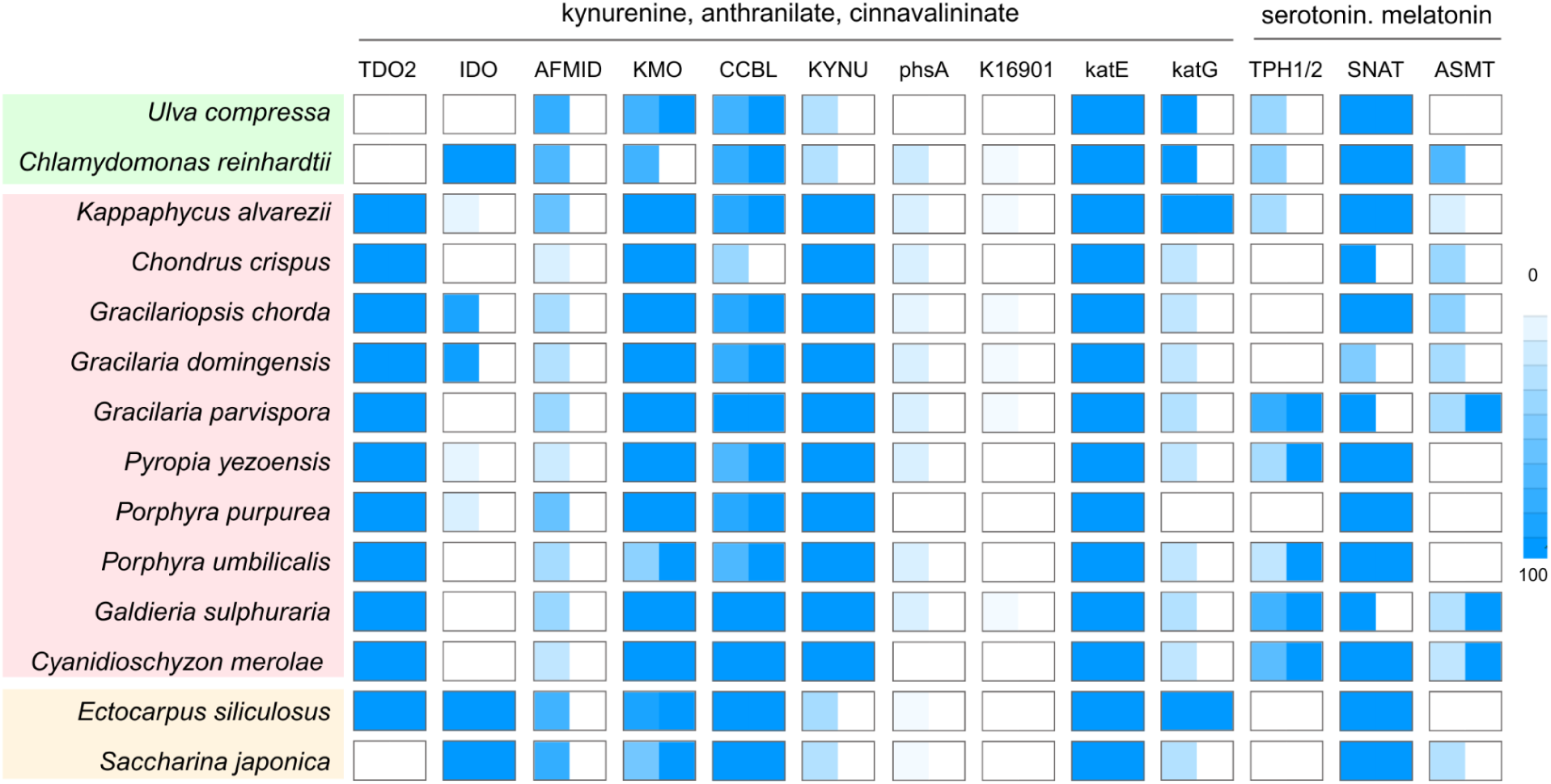
Heatmap comparison of algal orthologs for tryptophan-derived metabolite pathways, including kynurenine/anthranilate/cinnavalinate (TDO2, IDO, AFMID, KMO, CCBL, KYNU, phsA, K16901, katE, katG) and serotonin/melatonin biosynthesis (TPH1/2, SNAT, ASMT). Species are grouped by major algal lineages (chlorophytes, rhodophytes, glaucophytes, brown algae). Color intensity indicates KofamKOALA (left) and EggNOG (right) orthology scores (0–100).

Kynurenine is synthesized via the oxidative cleavage of tryptophan by either tryptophan dioxygenase (TDO) or indoleamine 2,3-dioxygenase (IDO), followed by conversion of the intermediate L-formylkynurenine to L-kynurenine by kynurenine formamidase (BNA7). L-kynurenine can then be metabolized into anthranilic acid (AA), 3-hydroxykynurenine, or kynurenic acid through the actions of kynureninase (KYNU), kynurenine 3-monooxygenase (KMO), and kynurenine-oxoglutarate transaminase (CCBL), respectively.

Expression analysis revealed that nearly all *K. alvarezii* genes in the kynurenine pathway were present and transcriptionally active. Specifically, AFMID, KMO, and CCBL were highly expressed and strongly light-responsive, suggesting photoregulated flux through this pathway (Fig. 8, Fig. S3). In contrast, KYNU expression was weaker, however still detectable in *K. alvarezii* and other rhodophytes, consistent with lineage-specific retention of this enzyme. Together, this broad expression and regulation pattern supports a central role for kynurenine metabolism in *K. alvarezii* physiology, possibly in modulating stress response or hormone crosstalk.

### Jasmonic Acid

Jasmonic acid (JA) is synthesized from α-linolenic acid (α-LeA, 18:3) via a multi-step pathway. In plants, the process begins in the chloroplast, where α-LeA is converted into OPDA by the sequential action of 13-lipoxygenase (LOX), allene oxide synthase (AOS), and allene oxide cyclase (AOC). OPDA is then transported to the peroxisome and reduced to OPC-8:0 by OPDA reductase (OPR). Subsequent steps involve activation by OPCL, and three rounds of β-oxidation via acyl-CoA oxidase (ACX), multifunctional protein (MFP), and 3-ketoacyl-CoA thiolase (KAT), ultimately producing JA. In the cytoplasm, JA is conjugated to isoleucine to form the bioactive JA-Ile.

Among the species examined, only phaeophytes encoded orthologs of both LOX and AOS, while all lacked AOC (Fig. 9). Downstream peroxisomal enzymes, including ACX, MFP, and fadA, were broadly present across lineages, except for OPCL, which was consistently absent. Expression analysis in *K. alvarezii* revealed that transcripts for ACX and MFP were among the most strongly represented within the pathway, consistent with the genomic heatmap. LOX and AOS orthologs showed no significant differential expression under the conditions tested. Notably, JA biosynthesis genes in *K. alvarezii* were downregulated under low light conditions (Fig. S7, Table S7).

**Fig. 9.**
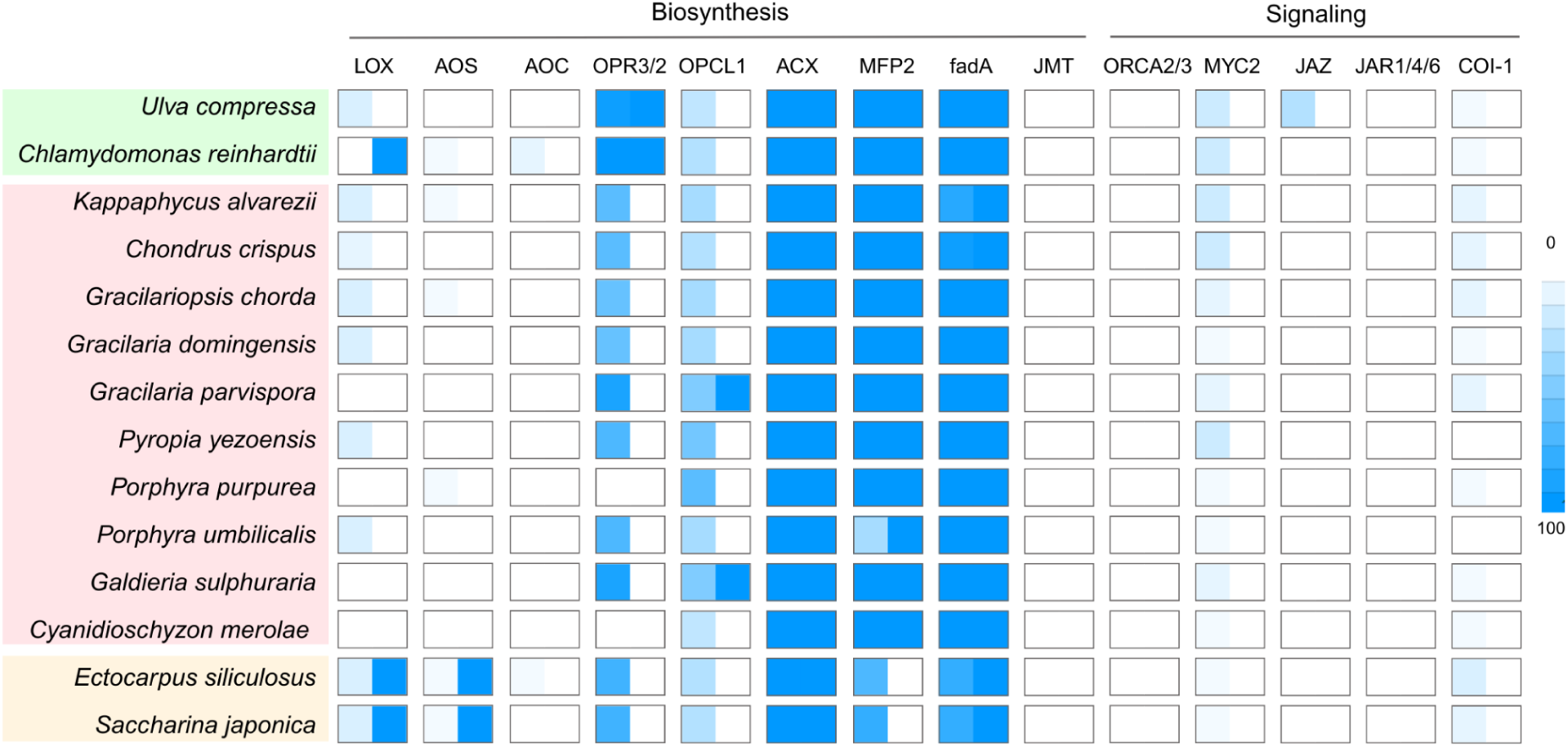
Heatmap comparison of algal orthologs for jasmonic acid (JA) biosynthesis (LOX, AOS, AOC, OPR3/2, OPCL1, ACX, MFP2, fadA, JMT) and signaling (ORCA2/3, MYC2, JAZ, JAR1/4/6, COI-1). Species are grouped by major algal lineages (chlorophytes, rhodophytes, glaucophytes, brown algae). Color intensity indicates KofamKOALA (left) and EggNOG (right) orthology scores (0–100)

### Salicyclic Acid

Salicyclic acid (SA) is synthesized from chorismate from two main pathways. In the isochorismate pathway, chorismate is converted to SA in two steps that are catalyzed by isochorismate synthase (ICS/pchA) and isochorismate pyruvate lyase (pchB). An alternative route, the phenylalanine ammonia-lyase (PAL) pathway, begins with the conversion of chorismate to phenylalanine by chorismate mutase (CM), followed by deamination to trans-cinnamic acid by PAL. This is further processed to benzoic acid via 3-hydroxyacyl-CoA dehydrogenase (AIM1) and ultimately hydroxylated to form SA, though the final enzyme remains uncharacterized (Lefevere et al, 2020; León et al, 1995).

Across algal species, pchA orthologs were universally detected, while pchB was identified in several taxa, including *K. alvarezii*, though generally at lower KofamKOALA stringencies. Of the signaling elements, only PR1, was detected in a few algal species examined (Fig. 10), underscoring the divergence of canonical SA signaling machinery.

**Fig. 10.**
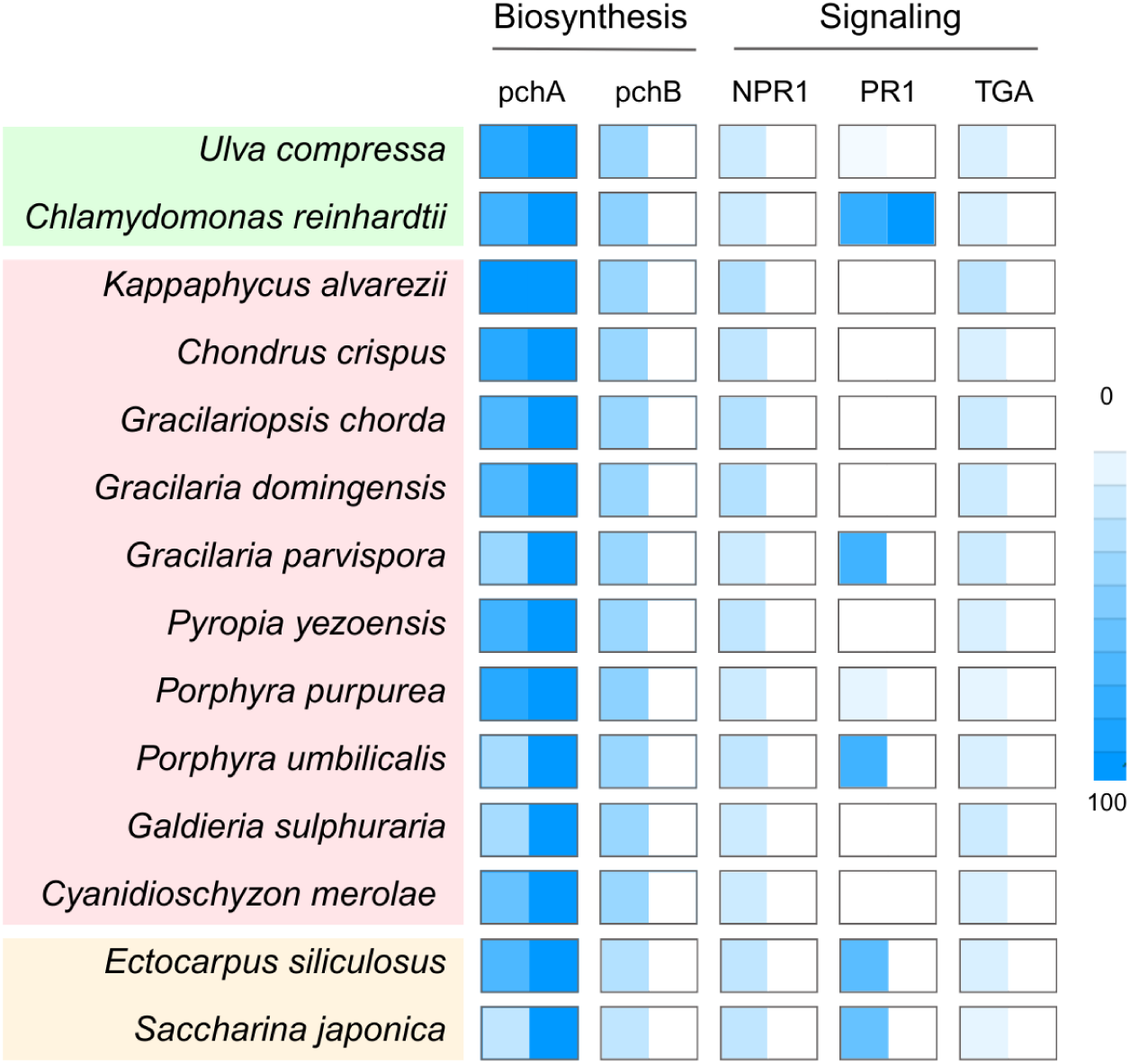
Heatmap comparison of algal orthologs for salicylic acid (SA) biosynthesis (pchA, pchB) and signaling (NPR1, PR1, TGA). Species are grouped by major algal lineages (chlorophytes, rhodophytes, glaucophytes, brown algae). Color intensity indicates KofamKOALA (left) and EggNOG (right) orthology scores (0–100).

In *K. alvarezii*, pchA expression was highest in medium light and lowest in low light, suggesting a light-responsive SA production (Fig. S8, Table S7).

### Strigolactones

Strigolactones (SLs) are synthesized from β-carotene through a series of enzymatic reactions. The isomerase DWARF27 (D27) first converts all-trans β-carotene into 9-cis-β-carotene, which is the cleaved by carotenoid cleavage dioxygenases (CCD7 and 8), forming carlactone, a key intermediate for strigolactone biosynthesis. Carlactone is further oxidized by cytochrome P450 enzymes, notably the More Auxiliary Growth 1 (MAX1) protein, into various canonical and non-canonical SLs. SL signaling is also mediated by the SCF E3 ubiquitin ligase complex: binding of SL to its receptor DWARF14 (D14) protomotes interaction with the F-box protein MAX2, forming the SCF^MAX2 complex, which targets SMXL (suppressor of MAX2-like repressors for degradation, allowing transcription of SL-responsive genes.

Across the species examined, D27 orthologs were widely detected and showed strong transcript representation, but CCD7 orthologs were completely absent and CCD8 orthologs were found in only four species (Fig. 11). This expression pattern suggests that while the initial isomerization and partial cleavage steps may be retained in algae, the complete canonical pathway is unlikely functional. Although the cytochrome P450 CYP711 family plays a key role in SL diversification in plants, specific orthologs were not identified, reflecting sequence divergence and variability in SL biosynthetic enzymes across taxa (Jia, Baz, and Al-Babili 2018). Similarly, no expression evidence was found for SL signaling components, (D14, MAX2, and SMXL) outside of charophytes, consistent with previous reports suggesting that canonical SL signaling may be restricted to streptophyte algae and land plants (Delaux et al, 2012; Stirk & Van Staden, 2014).

**Fig. 11.**
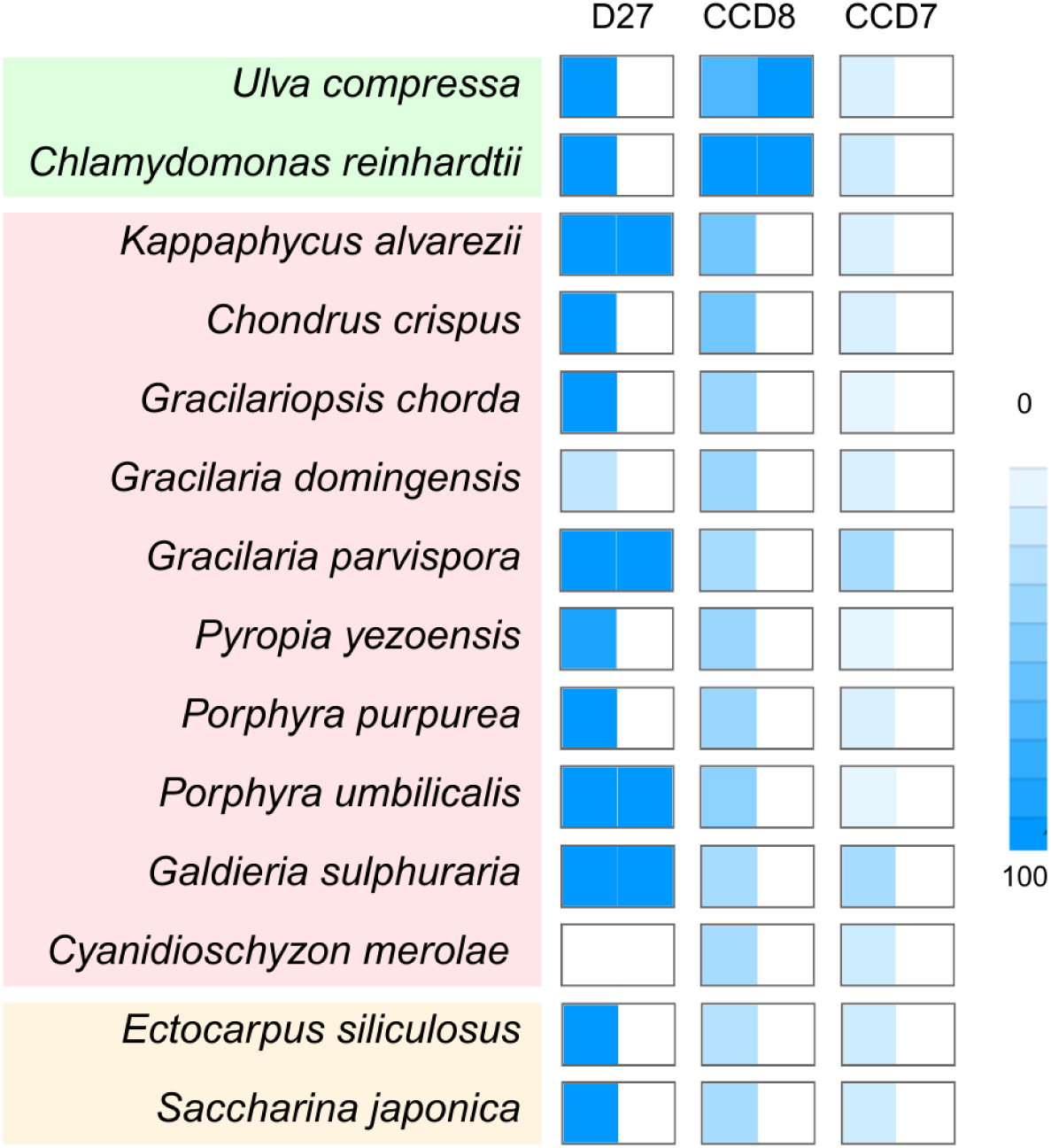
Heatmap comparison of algal orthologs for strigolactone (SL) biosynthesis (D27, CCD8, CCD7). Species are grouped by major algal lineages (chlorophytes, rhodophytes, glaucophytes, brown algae).

Color intensity indicates KofamKOALA (left) and EggNOG (right) orthology scores (0–100).

### Brassinosteroids

Brassinosteroids (BRs) are polyhydroxylated steroid hormones that play key roles in plant growth and stress response. They are classified into C27-, C28-, and C29-type BRs based on the number of carbon atoms, which correspond to their sterol precursors: cholesterol, campestrol, and sitosterol, respectively. All BR types share a common early biosynthetic pathway, which may proceed via the mevalonate (MVA) or non-MVA pathway, before diverging into distinct downstream conversions based on sterol substrate. BR synthesis involves C6 oxidation, catalyzed by enzymes including the 5α-reductase De-Etiolated-2 (DET2) and several cytochrome P450s (Bajguz et al, 2020; Kim et al, 2004; Ohnishi et al, 2006).

DET2 orthologs were identified in seven algal species, representing all major subdivisions (Fig. 12). Except for Chondrus crispus, all species had at least one cytochrome P450 gene associated with BR biosynthesis, most commonly CYP90C1/D1. In *K. alvarezii*, CYP90-related genes were detected with low-to-moderate annotation scores, but their expression levels were weak or absent, indicating limited transcriptional support for BR biosynthesis under the tested light conditions (Fig. S2).

**Fig. 12.**
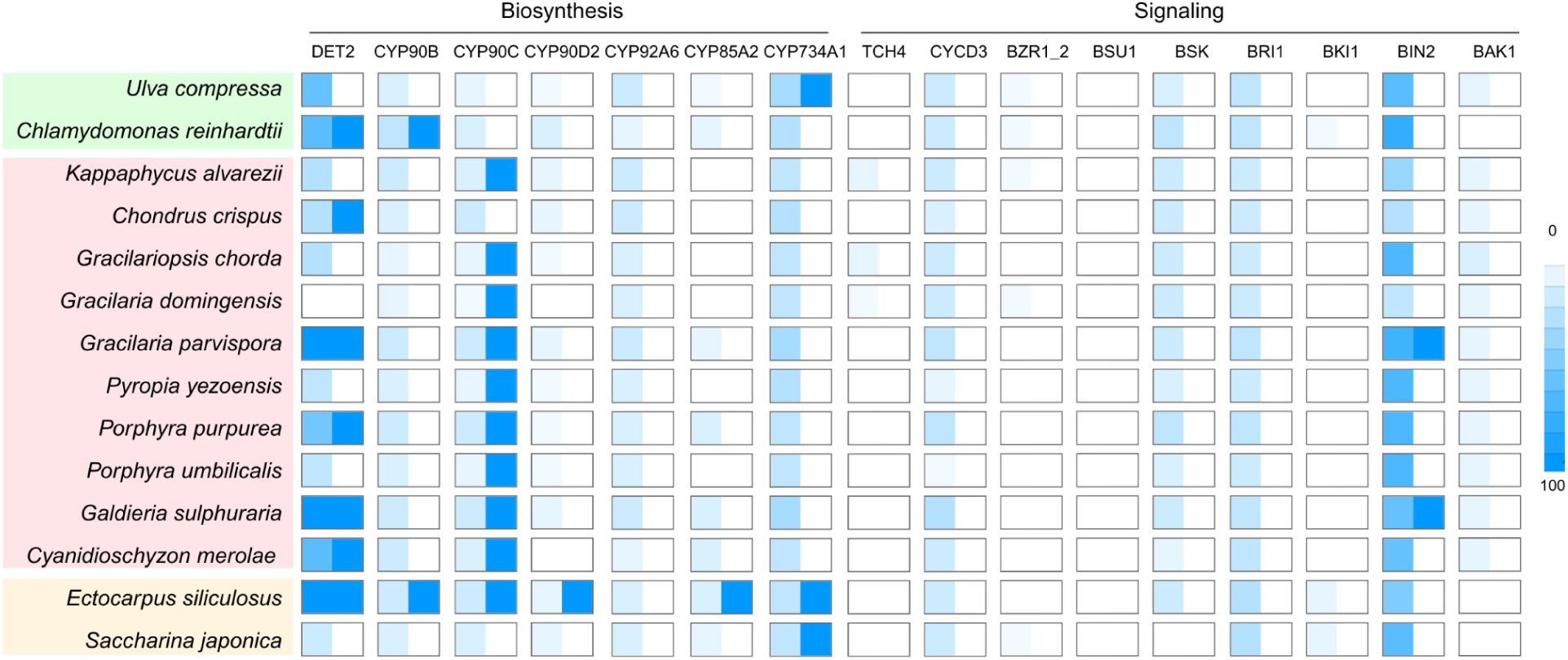
Heatmap comparison of algal orthologs for brassinosteroid (BR) biosynthesis (DET2, CYP90B, CYP90C, CYP90D2, CYP92A6, CYP85A2, CYP734A1) and signaling (TCH4, CYCD3, BZR1/2, BSU1, BSK, BRI1, BKI1, BIN2, BAK1). Species are grouped by major algal lineages (chlorophytes, rhodophytes, glaucophytes, brown algae). Color intensity indicates KofamKOALA (left) and EggNOG (right) orthology scores (0–100).

BR signaling in higher plants is initiated at the cell surface by the receptor kinase BRI1 and its co-receptor BAK1. Ligand binding induces dissociation of the inhibitor BRI1, enabling BRI1–BAK1 interaction and phosphorylation events that trigger a downstream signaling cascade. This cascade leads to inactivation of the negative regulator BIN2, allowing the transcription factors BES1 and BZR1 to accumulate in the nucleus and regulate BR-responsive genes (Belkhadir et al, 2006; Li et al, 2002; Tang et al, 2008; Zhao et al, 2002). BR signaling also intersects with other hormonal pathways, including ethylene, by stabilizing biosynthetic intermediates such as ACC and ACS.

Canonical BR signaling components were largely absent in algae. Only Chlamydomonas reinhardtii possessed a BAK1 ortholog, and although BIN2 homologs were present in most species, *K. alvarezii* lacked this component entirely. Expression data further confirmed these absences: while low-level transcription was observed for some signaling-related orthologs (e.g., BSK, BIN2 in other lineages), *K. alvarezii* showed no detectable expression of key BR signaling elements (Fig. 12).

## Discussion

Our genome-wide survey of phytohormone-related genes in *K. alvarezii* highlights a mosaic architecture in which conserved signaling modules are maintained despite incomplete or divergent biosynthetic pathways. This pattern points to an evolutionary compromise: while red algae lack many canonical plant enzymes, they appear to sustain hormonal responsiveness through alternative routes or microbial complementation. Such flexibility may be central to their ecological resilience and underlies the activity of *K. alvarezii-*derived biostimulants.

Auxin and tryptophan metabolites emerged as particularly well supported.The IAM pathway, long considered a minor route in plants, appears dominant in algae, with microbial-type iaaM genes strongly retained in *K. alvarezii*. This suggests horizontal gene transfer or co-evolution with bacteria, a scenario consistent with the well-documented microbial production of auxins in marine environments. The strong light responsiveness of auxin to photoadaptation, echoes observations in *Klebsormidium nitens* and other algae species. Importantly, the parallel induction of kynurenine inhibits auxin biosynthesis enzymes (He et al, 2011), and our results suggest a similar regulatory role may exist in algae, potentially coupling light perception to growth and morphology.

Cytokinin pathways in *K. alvarezii* reflect both conservation and divergence. While TRIT1 and CYP735A orthologs support the capacity to produce cZ and tZ, the absence of canonical IPT enzymes suggests a noncanonical initiation route. The light-dependent regulation of cytokinin genes aligns with their function in stress protection in plants (Cortleven et al, 2014) and with seasonal cycling in kelps (de Nys et al, 1990). The consistent detection of cytokinin-like activity in seaweed extracts (Stirk et al, 2003; 2009) may therefore be explained by both endogenous algal biosynthesis and microbial supplementation, highlighting a dual origin that could be leveraged to optimize biostimulant formulations.

Gibberellins represent a striking case of divergence in *K. alvarezii*. The absence of CPS/KS suggests that algae have either lost or replaced canonical upstream enzymes, yet the conservation of GA oxidases and their strong affinity for GA4 and GA3 implies downstream activity is functional. Unlike the other hormones analyzed, GA underwent further structural modeling and ligand docking because conventional annotation failed to recover canonical biosynthetic genes, despite previous reports of endogenous GA production in algae (Prasad et al, 2010; Yalçin et al, 2019). This partial retention of GA activity has been reported to stimulate growth and branching in some algal species (Burkiewicz, 1987; Borowczak et al, 2008; Park et al, 2013) but inhibit or produce negligible effects in others (Garcia-Jimenez et al, 1998; Stirk et al, 2019). The presence of GA2ox-like proteins, supported by structural conservation, docking with GA8 and GA34, and strong expression under high light conditions, suggests that GA inactivation is a central regulatory mechanism in *K. alvarezii*. In higher plants, GA2ox activity was pivotal to semi-dwarf crop traits during the Green Revolution (Hedden et al, 2003; Sasaki et al, 2002). In algae, similar regulatory “brakes” may allow growth to be fine-tuned in dynamic environments. Together, these results provide the first structural and functional evidence for GA metabolism in *K. alvarezii,* pointing to a balance between biosynthesis and inactivation that may directly influence the biostimulant efficacy of its extracts.

Abscisic acid findings reinforce the importance of signaling over biosynthesis. While most upstream enzymes were absent, the presence of ABA3 and multiple signaling orthologs (PP2C, SnRK2, ABF), combined with strong light-responsive expression, suggests that *K. alvarezii* is capable of perceiving and responding to ABA even if synthesis depends on alternative routes or microbial input. This resonates with reports of microbial ABA production in marine bacteria (Forchetti et al, 2007; Shukla et al, 2012) and suggests that algae may outsource biosynthesis while retaining endogenous signaling capacity. The conservation of stress-responsive transcriptional control highlights ABA as a key integrator of algal acclimation, consistent with its roles in oxidative stress, salinity, and desiccation tolerance across diverse taxa (Guajardo et al, 2016; Sun et al, 2022; Yoshida, 2006).

Ethylene and ACC signaling similarly illustrate this modular architecture. The presence of ACS but not ACO indicates an incomplete pathway, yet strong light induction of ACS suggests active production of ACC. Recent work shows that ACC itself functions as a signaling molecule independent of ethylene (Uji & Mizuta, 2022; Yanagisawa et al, 2019), a finding that fits neatly with our genomic and expression data. Rather than being viewed as a truncated pathway, the algal ethylene network may represent a bifurcated system, capable of sensing both ethylene and ACC under different environmental contexts.

Jasmonic acid and salicylic acid were the least conserved, yet their functional relevance is supported by expression data and physiological studies. In *K. alvarezii*, downregulation of JA-related genes under low light and upregulation of SA biosynthesis under medium light suggest dynamic partitioning of stress responses between these two hormones. While canonical signaling modules are absent, algae consistently respond to exogenous JA and SA (Bouarab et al, 2004; Hu et al, 2021; Zhou et al, 2010), indicating the existence of divergent perception systems. Such findings raise the possibility that JA- and SA-like molecules in algae are not strictly homologous to plant hormones but still trigger analogous defense and stress pathways.

Strigolactones and brassinosteroids were the most reduced. SL biosynthesis appeared limited to early steps, and BR pathways, while retaining DET2 and CYP90, lacked signaling elements such as BRI1 and BAK1. The very weak or absent expression of BR-related genes in *K. alvarezii* suggests that, even if biochemically possible, these pathways are unlikely to operate in a canonical form. Nevertheless, SL- or BR-like activity has been reported in seaweed extracts (Arioli et al, 2015; Bajguz, 2000), suggesting either microbial origin or alternative biosynthetic solutions (Table 1).

**Table 1:**
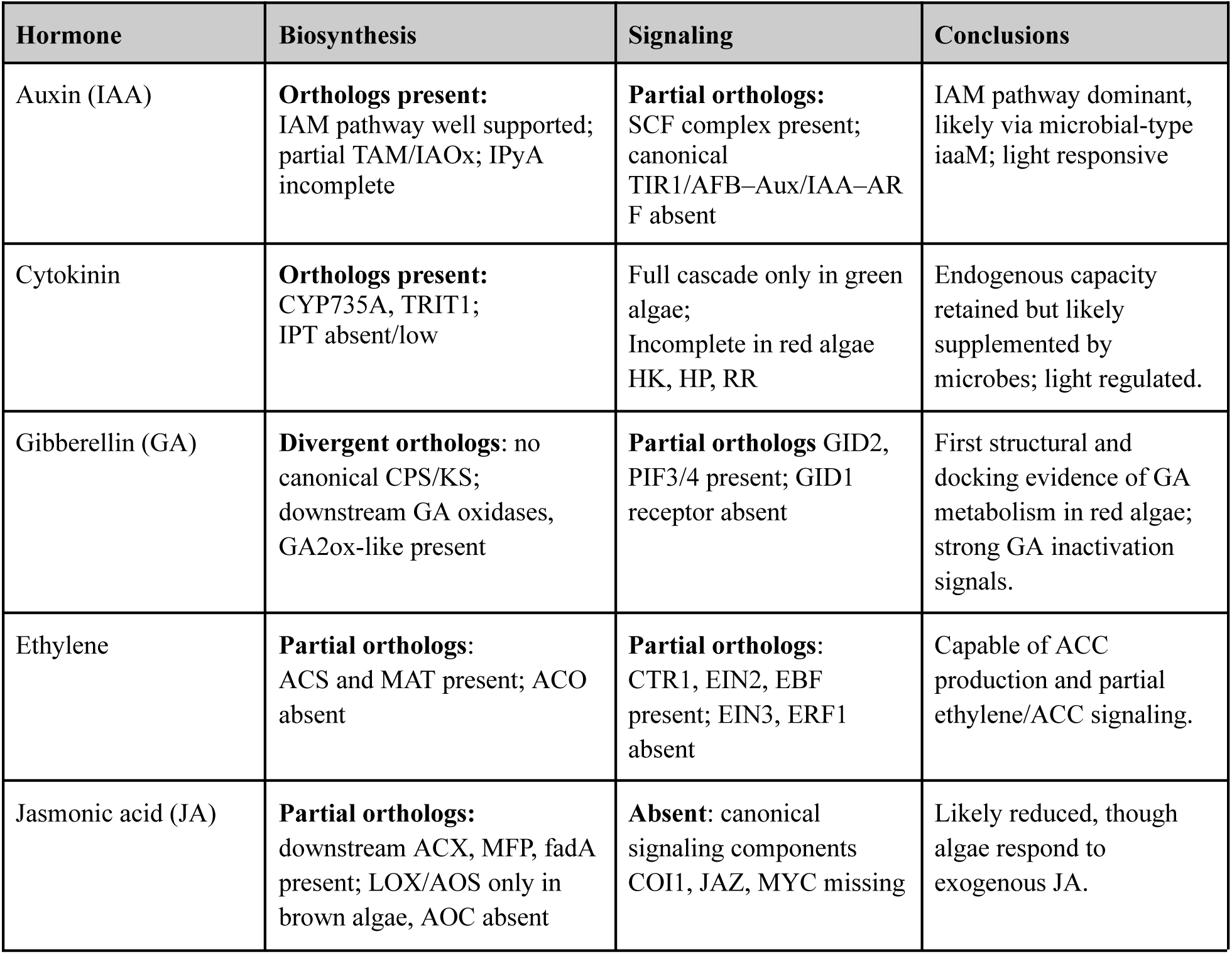
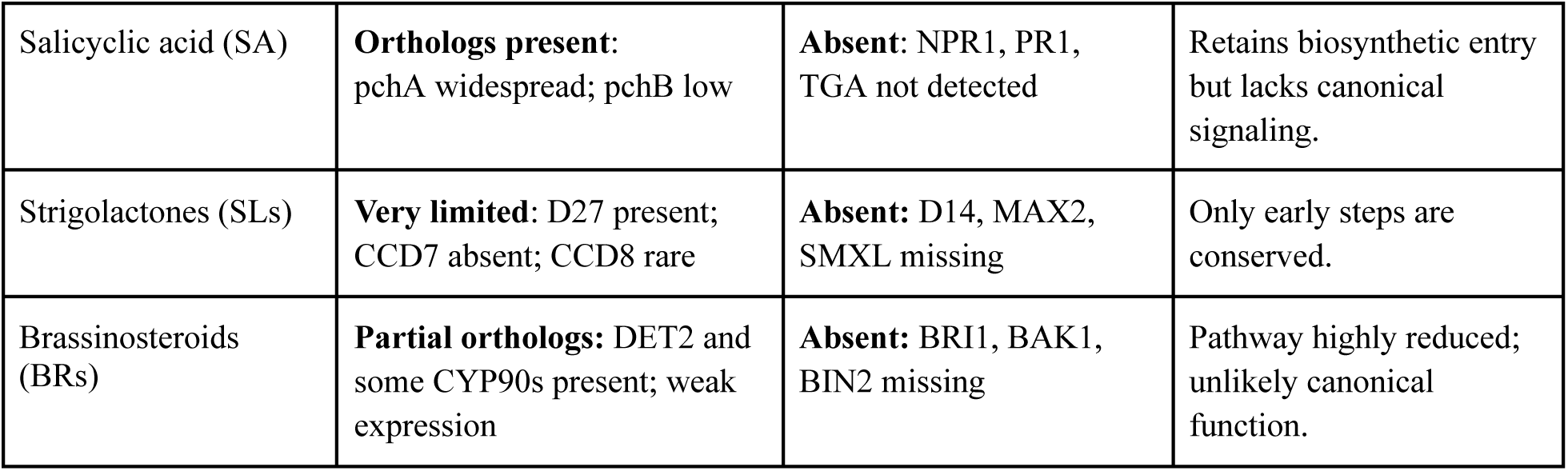
Summary of Phytohormone Biosynthesis and Signaling Components in algae.

| Hormone | Biosynthesis | Signaling | Conclusions |
| --- | --- | --- | --- |
| Auxin (IAA) | **Orthologs present:**<br>IAM pathway well supported;<br>partial TAM/IAOx; IPyA incomplete | **Partial orthologs:**<br>SCF complex present;<br>canonical<br>TIR1/AFB–Aux/IAA–AR<br>F absent | IAM pathway dominant,<br>likely via microbial-type<br>iaaM; light responsive |
| Cytokinin | **Orthologs present:**<br>CYP735A, TRIT1;<br>IPT absent/low | Full cascade only in green<br>algae;<br>Incomplete in red algae<br>HK, HP, RR | Endogenous capacity<br>retained but likely<br>supplemented by<br>microbes; light regulated. |
| Gibberellin (GA) | **Divergent orthologs:** no<br>canonical CPS/KS;<br>downstream GA oxidases,<br>GA2ox-like present | **Partial orthologs** GID2,<br>PIF3/4 present; GID1<br>receptor absent | First structural and<br>docking evidence of GA<br>metabolism in red algae;<br>strong GA inactivation<br>signals. |
| Ethylene | **Partial orthologs:**<br>ACS and MAT present; ACO<br>absent | **Partial orthologs:**<br>CTR1, EIN2, EBF<br>present; EIN3, ERF1<br>absent | Capable of ACC<br>production and partial<br>ethylene/ACC signaling. |
| Jasmonic acid (JA) | **Partial orthologs:**<br>downstream ACX, MFP, fadA<br>present; LOX/AOS only in<br>brown algae, AOC absent | **Absent:** canonical<br>signaling components<br>COI1, JAZ, MYC missing | Likely reduced, though<br>algae respond to<br>exogenous JA. |
| Salicylic acid (SA) | **Orthologs present:** pchA widespread; pchB low | **Absent:** NPR1, PR1, TGA not detected | Retains biosynthetic entry but lacks canonical signaling. |
| Strigolactones (SLs) | **Very limited:** D27 present; CCD7 absent; CCD8 rare | **Absent:** D14, MAX2, SMXL missing | Only early steps are conserved. |
| Brassinosteroids (BRs) | **Partial orthologs:** DET2 and some CYP90s present; weak expression | **Absent:** BRI1, BAK1, BIN2 missing | Pathway highly reduced; unlikely canonical function. |

Across all hormones, a common theme emerges: biosynthesis is often incomplete or divergent, but signaling modules are selectively retained and transcriptionally responsive to light stress. This pattern suggests that algae rely on a combination of endogenous enzymes, microbial partners, and alternative perception mechanisms to achieve hormonal regulation. For *K. alvarezii*, these findings provide a mechanistic explanation for the observed bioactivity of its extracts and underscore the importance of environmental cues such as light in shaping hormone-related responses.

This study therefore establishes a molecular framework for understanding the endogenous hormone biology of *K. alvarezii* and its contribution to the biostimulant activity of its extracts. Our results demonstrate that, while canonical hormone signaling architectures are incomplete in red algae, orthologs were detected for all nine major phytohormone classes. Structural modeling and ligand docking provided functional support for candidate enzymes such as GGPPS- and GA oxidase–like proteins, while transcriptomic profiling revealed light-responsive regulation across auxin, cytokinin, ethylene, salicylic acid, and ABA pathways. These findings highlight both the conservation and environmental responsiveness of algal hormone systems, underscoring their likely roles in growth regulation and stress tolerance.

Future research should now prioritize biochemical validation of key enzymes, quantitative hormone profiling under controlled environmental conditions, and functional assays for divergent candidates such as GTG1/GTG2 in ABA perception and PILS transporters in auxin transport. Investigation of microbial partners that could complement incomplete biosynthetic pathways is also critical, as symbionts may provide missing enzymatic capacity (Mikami et al, 2016; Wichard et al, 2015).

From an applied perspective, linking endogenous hormone profiles to biostimulant efficacy provides a clear path forward. Reports that GA removal from *K. alvarezii* extracts enhances crop performance (Ghosh et al, 2014; Mondal et al, 2015) highlight the value of tailoring hormonal composition through strain selection, cultivation practices, or extract processing. Establishing standardized extraction protocols and benchmarks for hormone quantification will be essential to ensure consistency and efficacy. By integrating molecular insights, microbial ecology, and applied testing, *K. alvarezii* can be advanced as a sustainable and scientifically optimized biostimulant, strengthening its role in agriculture and aquaculture.

## Data Availability

All data supporting the findings of this study are available in the supplementary repository associated with this article. Supplementary Tables S1–S9b are deposited in GitHub and archived in Zenodo to ensure long-term accessibility (https://doi.org/10.5281/zenodo.17139152). A detailed README file describing the content and structure of each table is included in the repository. Full data will be available through NCBI upon acceptance.

## Supporting information

All_supplementary_tabs_figs

## Acknowledgements

This study was supported by the research fund from the United Kingdom Research and Innovation–Global Challenges Research Fund (UKRI-GCRF) ‘GlobalSeaweedSTAR’ Programme GSS/RF/015 (BB/P027806/1) awarded to MYR. Publication cost support provided by USC SBIR SVN.

## Author Contribution Statement

**Bea A. Crisostomo:** Methodology, Formal analysis, Investigation, Data Curation, Writing - Original Draft, Writing - Review & Editing, Visualization. **Leah Ferger:** Investigation, Data curation, Writing-Reviewing and Editing, Visualization. **April Mae Tabona-Nabor:** Investigation, Data Curation, Writing - Review & Editing. **Alexandros Aramos:** Methodology, Software, Formal analysis, Data Curation, Writing - Review & Editing, Visualization. **Gabriel Schweizer:** Formal analysis, Methodology, Data curation, Software. **Sergey Nuzhdin:** Funding acquisition, Supervision, Writing - Review & Editing. **Arturo O. Lluisma:** Resources, Writing - Review & Editing, Supervision, Funding acquisition. **Michael Y. Roleda:** Resources, Writing - Review & Editing, Supervision, Project administration, Funding acquisition. **Scott Fahrenkrug:** Conceptualization, Methodology, Formal analysis, Data Curation, Writing - Review & Editing, Supervision, Project administration, Funding acquisition

