## Supplementary material for "Phytohormones in *Kappaphycus alvarezii:* Genomic Insights, Activation Thresholds, and Implications for Seaweed-derived Biostimulants": All_supplementary_tabs_figs

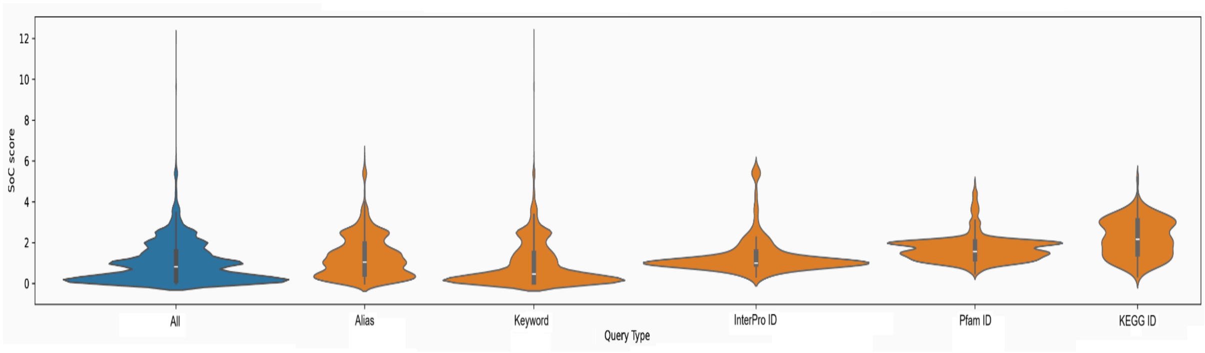


**Fig. S1** Violin plots of SoC scores across query term classes (All, Keyword, InterPro ID, Alias, KEGG ID, Pfam). White dots represent medians, boxes represent interquartile ranges, and density shapes show score distributions. Descriptions of each query group are provided in Table S5.

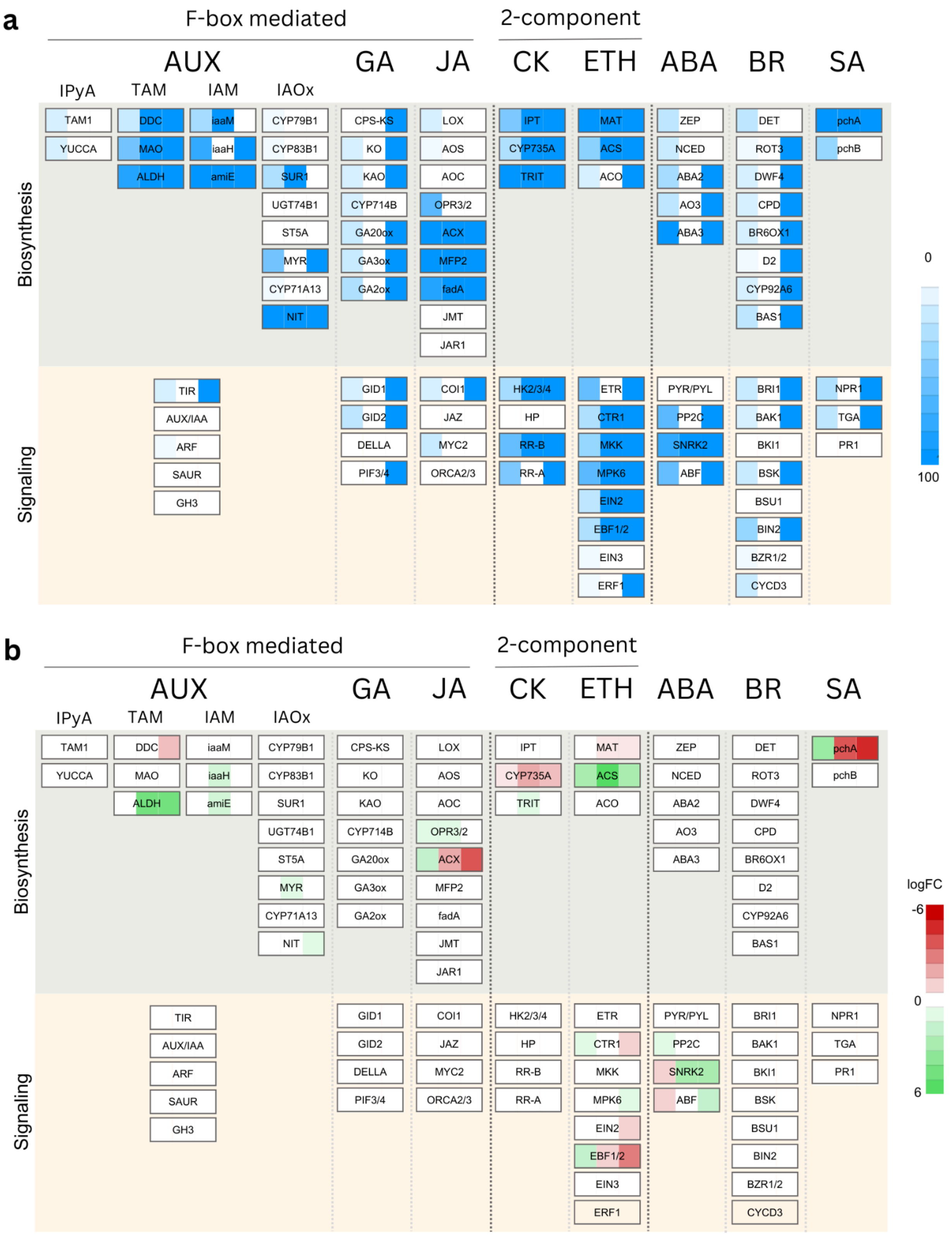


**Fig. S2** Distribution of algal orthologs for hormone biosynthesis and signaling pathways. (a) Presence/absence heatmap with orthology scores (0–100), highlighting conserved components across major phytohormone pathways.

The panels represent the following: KofamKOALA (left) scores, the eggNOG-mapper scores (middle) and the KaGE search results (right). (b) Differential expression of orthologs shown as log fold-change (logFC), indicating up- or down-regulation of candidate genes. The panels represent the following contrasts: High vs Low light (left), High vs Medium light (middle), Low vs Medium light (right). Pathways include auxin (AUX), gibberellin (GA), jasmonic acid (JA), cytokinin (CK), ethylene (ETH), abscisic acid (ABA), brassinosteroid (BR), and salicylic acid (SA).

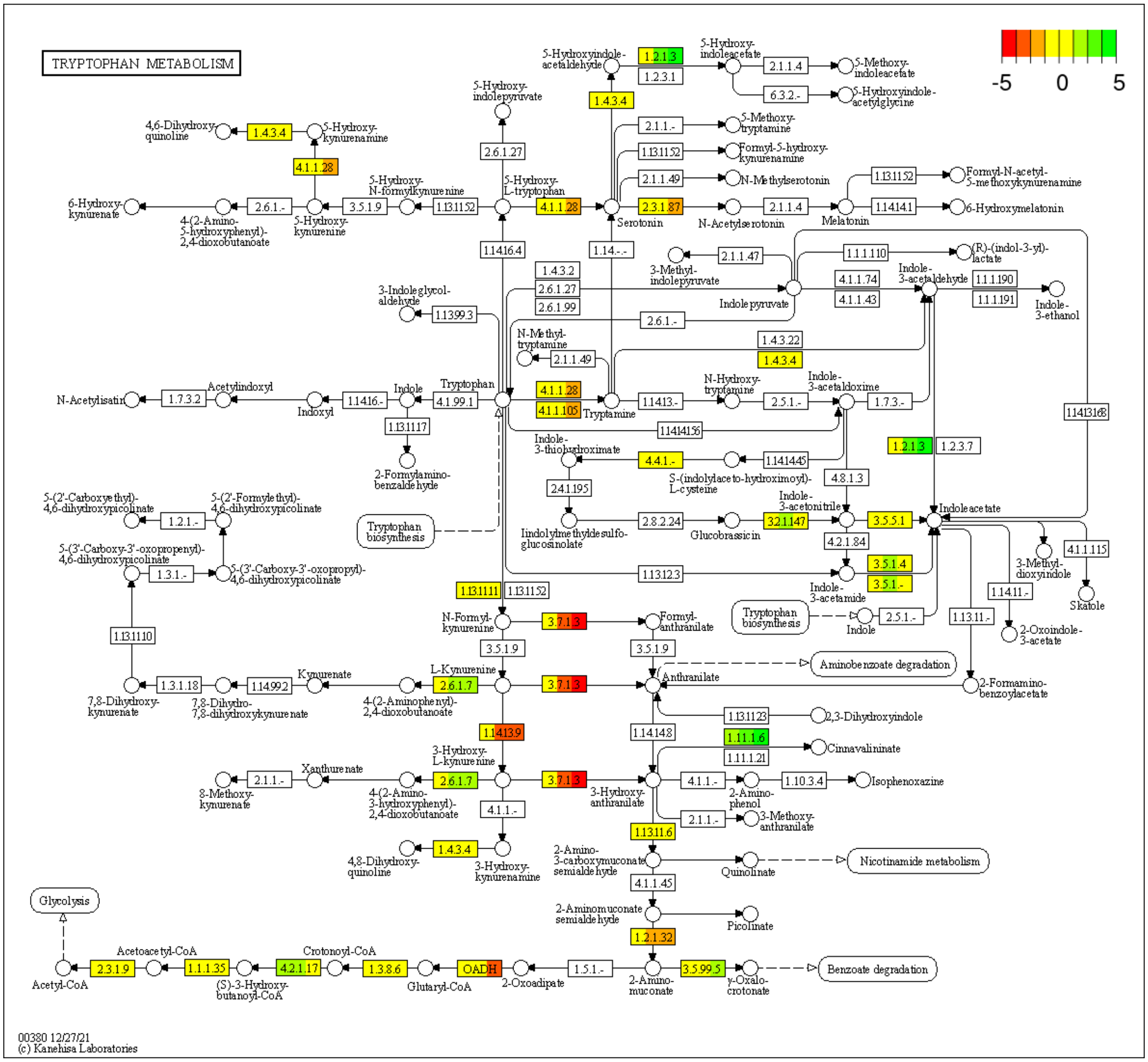


**Fig. S3** Effect of light intensity on the biosynthesis of auxin and other tryptophan-derived metabolites in *Kappaphycus alvarezii*. Genes in boxes are colored by log fold-change (logFC) under High vs. Low light (left), High vs. Medium light (middle), and Low vs. Medium light (right).

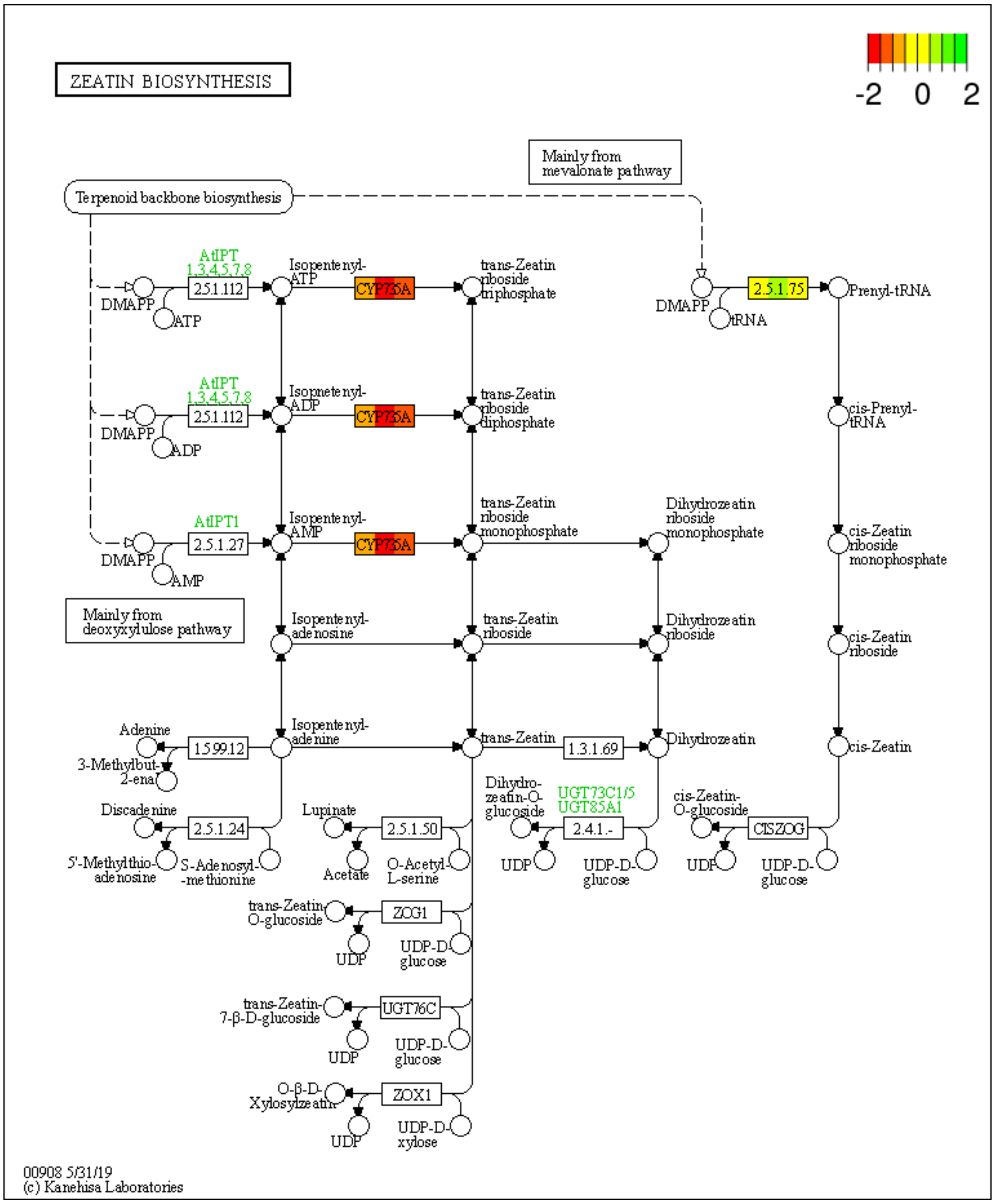


**Fig. S4** Effect of light intensity on cytokinin biosynthesis in *Kappaphycus alvarezii*. Genes in boxes are colored by log fold-change (logFC) under High vs. Low light (left), High vs. Medium light (middle), and Low vs. Medium light (right)

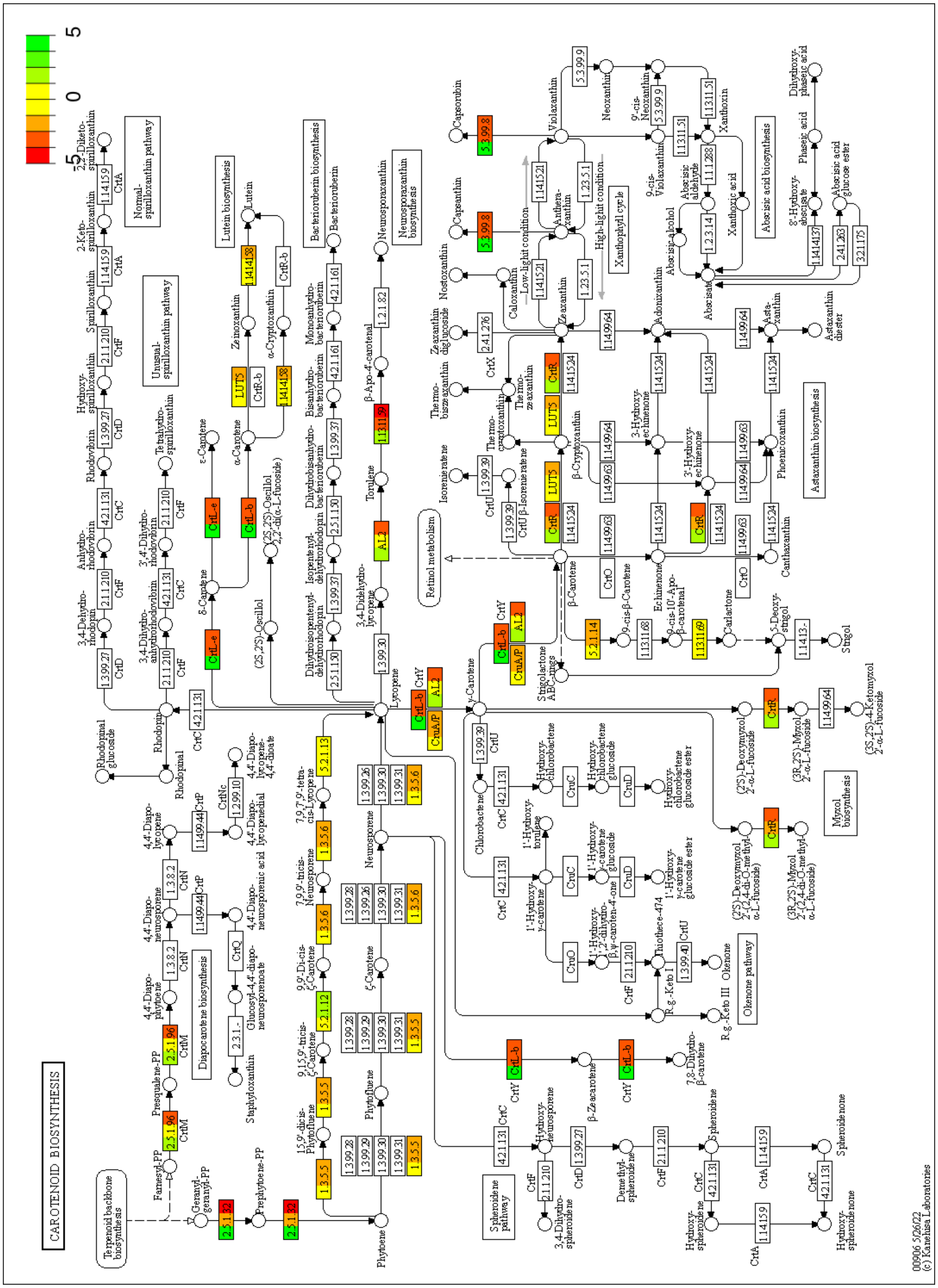


**Fig. S5** Effect of light intensity on carotenoid biosynthesis in *Kappaphycus alvarezii*. Genes in boxes are colored by log fold-change (logFC) under High vs. Low light (left), High vs. Medium light (middle), and Low vs. Medium light (right)

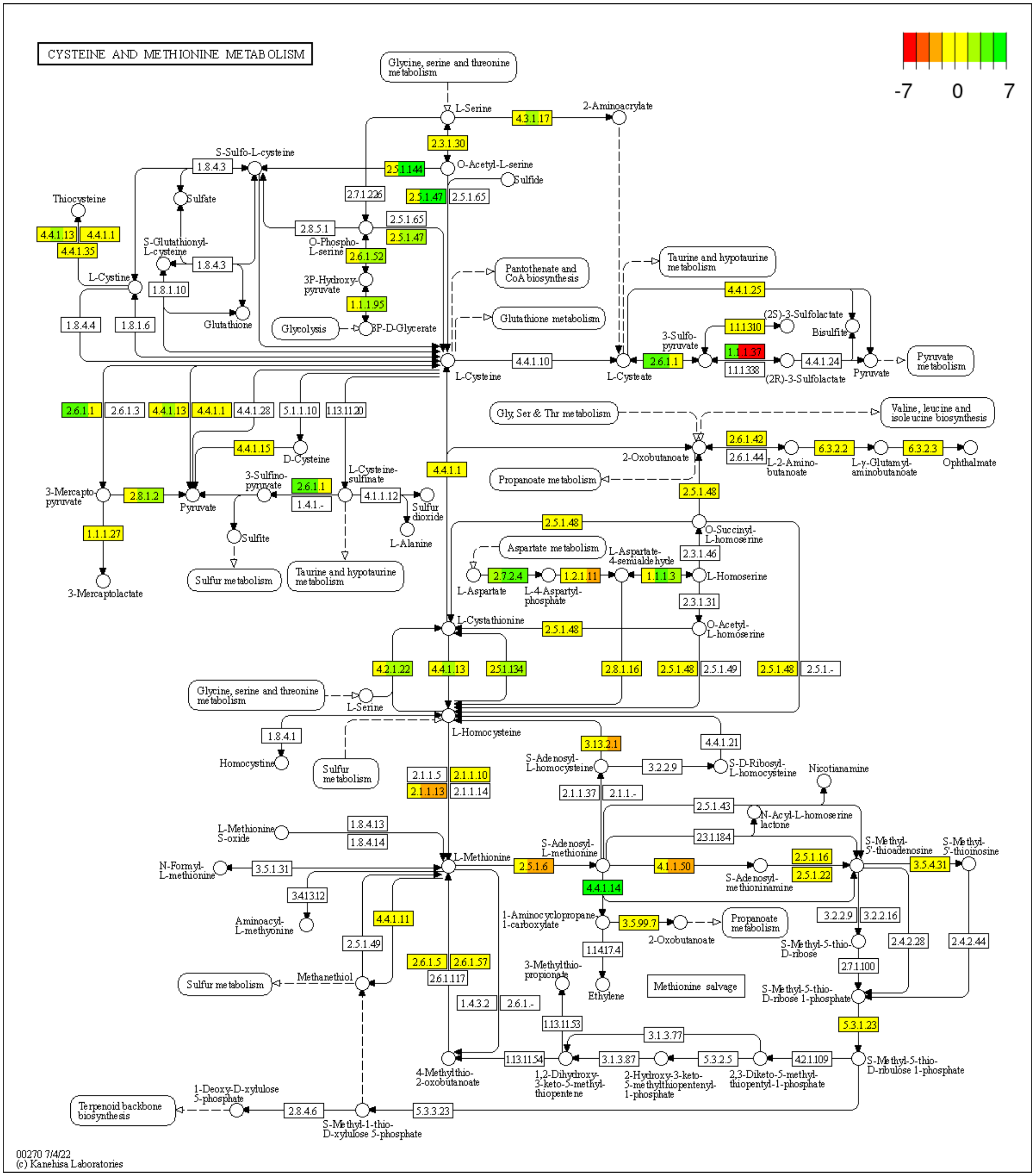


**Fig. S6** Effect of light intensity on cysteine and methionine metabolism in *Kappaphycus alvarezii*. Genes in boxes are colored by log fold-change (logFC) under High vs. Low light (left), High vs. Medium light (middle), and Low vs. Medium light (right)

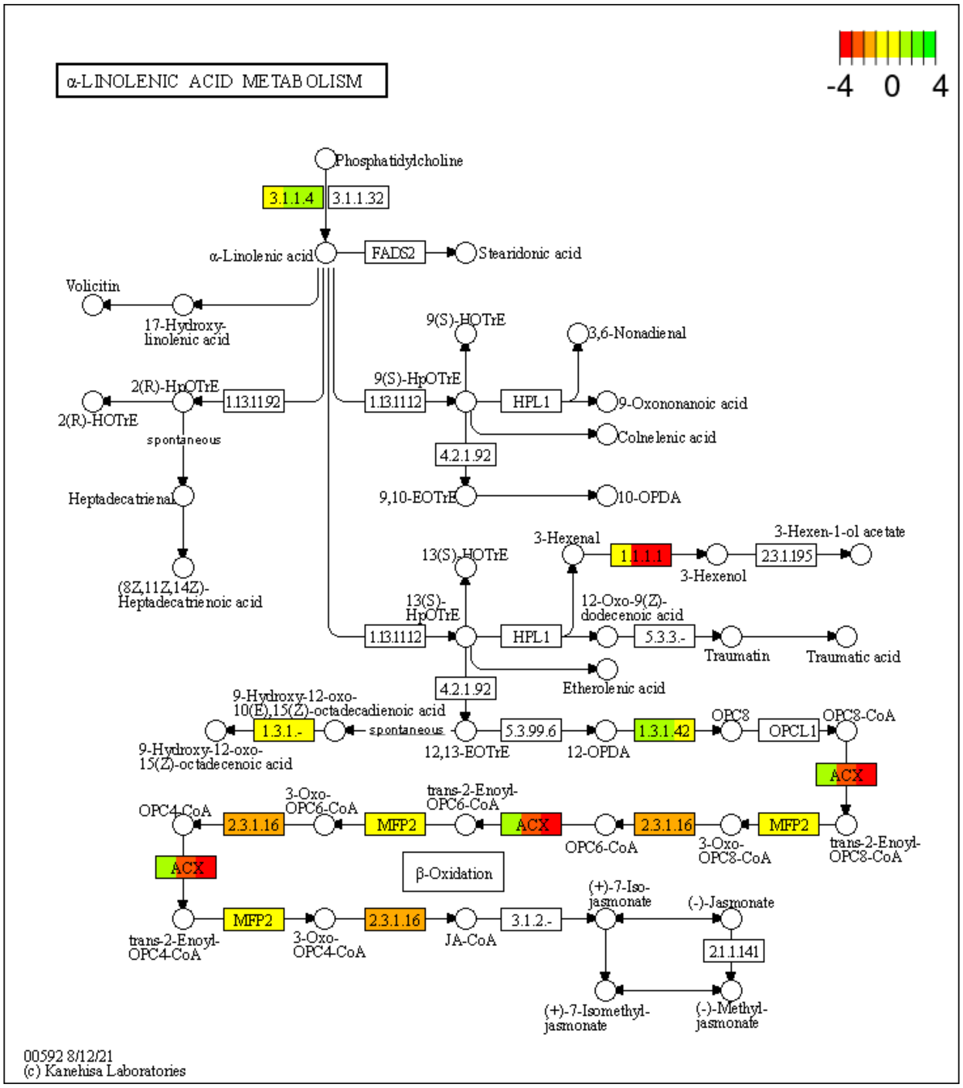


**Fig. S7** Effect of light intensity on jasmonic acid biosynthesis in *Kappaphycus alvarezii*. Genes in boxes are colored by log fold-change (logFC) under High vs. Low light (left), High vs. Medium light (middle), and Low vs. Medium light (right).

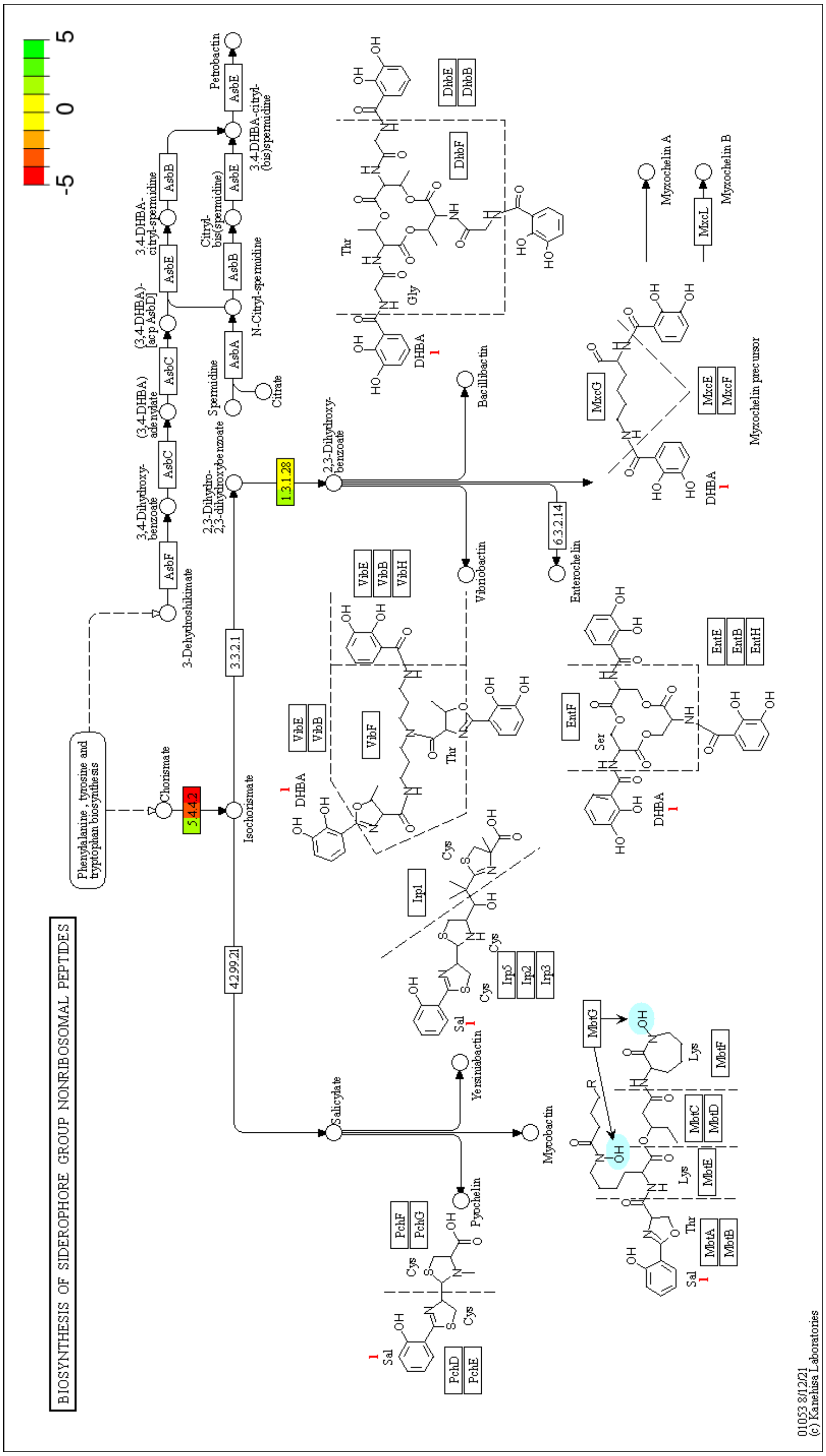


**Fig. S8** Effect of light intensity on salicylic acid biosynthesis in *Kappaphycus alvarezii*. Genes in boxes are colored by log fold-change (logFC) under High vs. Low light (left), High vs. Medium light (middle), and Low vs. Medium light (right)

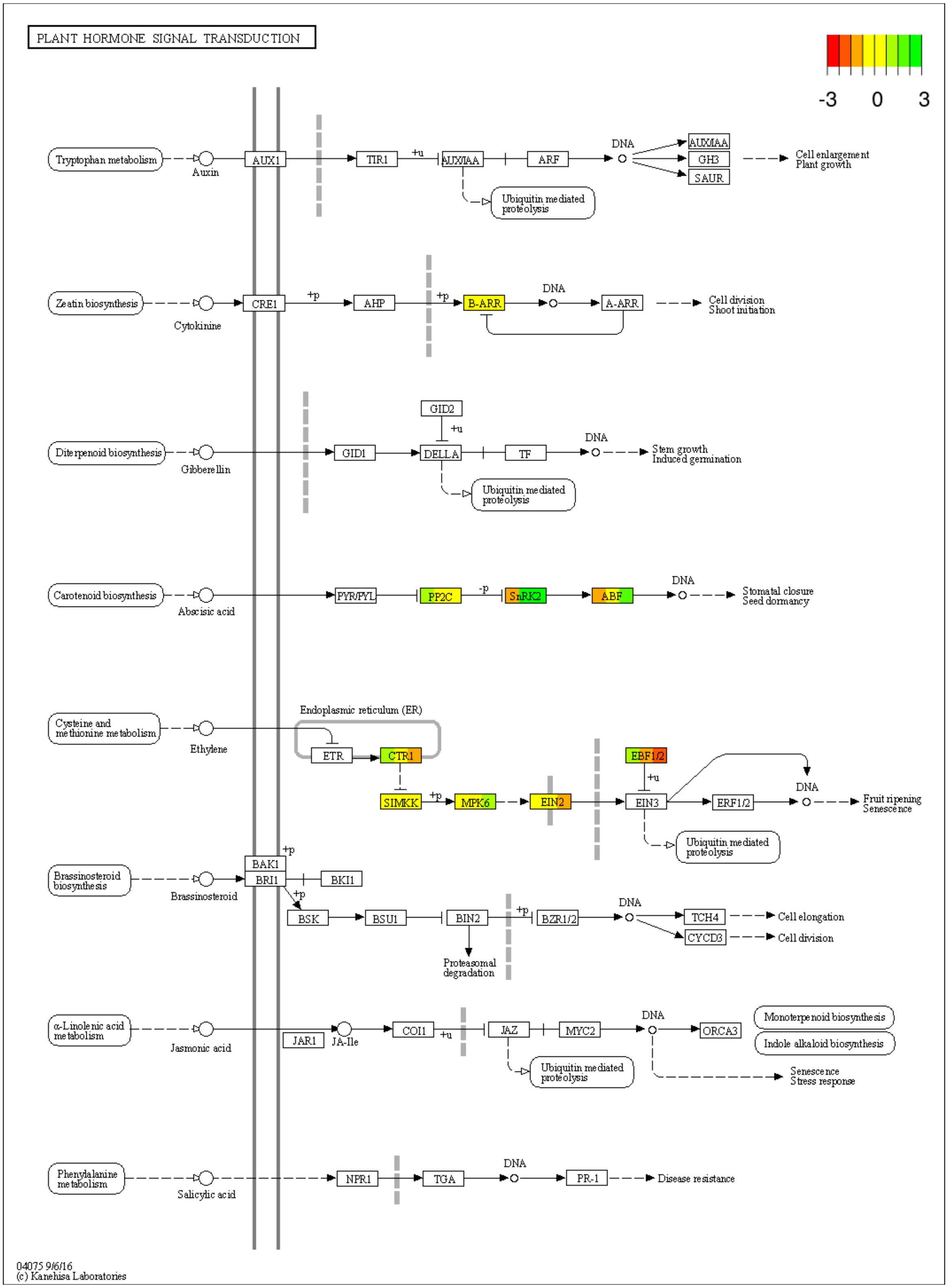


**Fig. S9** Effect of light intensity on phytohormone signaling in *Kappaphycus alvarezii*. Genes in boxes are colored by log fold-change (logFC) under High vs. Low light (left), High vs. Medium light (middle), and Low vs. Medium light (right)

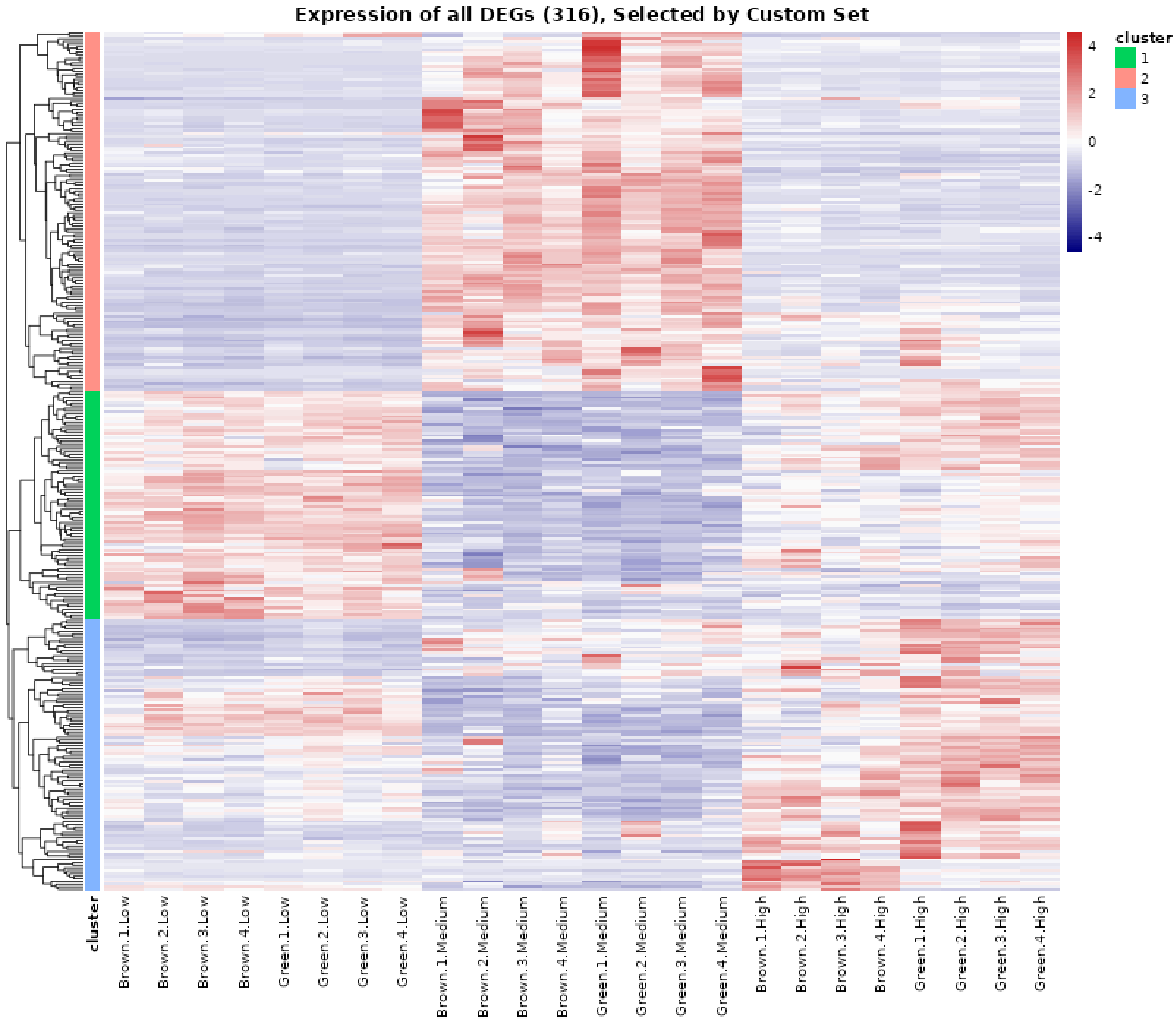


**Fig. S10** Expression of all differentially expressed phytohormone-related genes in *Kappaphycus alvarezii*. Genes with FDR ≤ 0.05 and a |logFC| ≥ 1 were considered differentially expressed. Clusters were assigned using the K.A. RShiny v2.3 tool
